# A reproducible human brain tissue model to study physiological and disease-associated microglia phenotypes

**DOI:** 10.64898/2026.01.20.699077

**Authors:** Julien Klimmt, Carolina Cardoso Gonçalves, Jessica Valentina Montgomery, Stephan A. Müller, Merle Bublitz, Severin Filser, Lars Paeger, Brigitte Nuscher, Angelika Dannert, Sigrun Roeber, Veronica Pravata, Martina Schifferer, Joshua J. Shrouder, Nathalie Schulz, Judit González Gallego, Silvia Cappello, Thomas Misgeld, Nikolaus Plesnila, Eduardo Beltrán, Jochen Herms, Elena De Domenico, Marc D. Beyer, Joachim L. Schultze, Christian Haass, Stefan F. Lichtenthaler, Caterina Carraro, Dominik Paquet

**Affiliations:** Institute for Stroke and Dementia Research (ISD), LMU University Hospital, LMU Munich, Munich, Germany; Graduate School of Systemic Neuroscience (GSN), LMU Munich, Munich, Germany; Systems Medicine, Deutsches Zentrum für Neurodegenerative Erkrankungen (DZNE) e.V., Bonn, Germany; German Center for Neurodegenerative Diseases (DZNE) Munich, Munich, Germany; Neuroproteomics, School of Medicine and Health, TUM University Hospital, Technical University of Munich, Munich, Germany; Center for Neuropathology and Prion Research, LMU Munich, Munich, Germany; Metabolic Biochemistry, Biomedical Center (BMC), Faculty of Medicine, LMU Munich, Munich, Germany; Department of Physiological Genomics, Biomedical Center (BMC), Faculty of Medicine, LMU Munich, Munich, Germany; Munich Cluster for Systems Neurology (SyNergy), Munich, Germany; Institute of Neuronal Cell Biology, Technical University of Munich, Munich, Germany; Institute of Clinical Neuroimmunology, LMU University Hospital, LMU Munich, Munich, Germany; Biomedical Center (BMC), Faculty of Medicine, LMU Munich, Munich, Germany; PRECISE Platform for Genomics and Epigenomics, Deutsches Zentrum für Neurodegenerative Erkrankungen (DZNE) e.V. and University of Bonn and West German Genome Center, Bonn, Germany; Genomics and Immunoregulation, Life & Medical Sciences (LIMES) Institute, University of Bonn, Bonn, Germany

## Abstract

Stem-cell-based *in vitro* models offer promising potential to elucidate human brain cell functions and interactions under physiological and pathological conditions. However, harnessing this potential is impaired by low reproducibility, maturity, or cell-type diversity of existing models. Especially, prolonged incorporation of mature microglia and studies of neuroinflammation have proven challenging. Here, we developed a 3D cortical brain tissue model (3BTM) containing neurons, astrocytes, and microglia with high reproducibility, maturity, and viability. 3BTMs show morphological, functional, and proteomic maturation of all cell types, leading to high similarity to their *in vivo* counterparts. Incorporated microglia survive for over 6 months and display mature morphology, functions, and gene expression. Importantly, when engineered to model Alzheimer’s disease pathology, 3BTMs recapitulate key disease hallmarks including amyloid deposition, increased phospho-Tau levels, and neuroinflammation, with microglia shifting their transcriptional landscape to disease-relevant signatures. Together, our model offers unprecedented possibilities for studying physiological and pathological states of human brain tissue and translational applications.

## Introduction

Investigating the physiological and disease-related interplay of human neurons and glia requires experimentally accessible and well-characterized model systems. Recent progress in the field of stem cell research has allowed generating multiple brain cell types from induced pluripotent stem cells (iPSC)^1^ and studying their interactions in co-cultures^2^. Moreover, these human *in vitro* systems enable the investigation of pathomechanisms and pathogenesis in disease-relevant cell types^3,4^. Although two-dimensional (2D) models containing neurons and glia co-cultured in a dish allow relatively simple model generation^5^, brain cells in 2D culture often lack physiological morphology, maturation, homeostatic behaviors, and intercellular crosstalk found in complex tissues^6–8^. 3-dimensional (3D) *in vitro* models offer the possibility to study cell-cell and cell-matrix interactions in a more physiological environment compared to 2D cultures. Furthermore, in Alzheimer’s disease (AD) models, they enable investigation of extracellular pathologies, such as the accumulation and aggregation of amyloid-β (Aβ) peptides, in addition to intracellular events such as tau phosphorylation and misfolding.

Current human iPSC-based 3D models include hydrogel-based systems, spheroid-like cultures, and neural organoids (NO). Hydrogel-based systems are generated by embedding dissociated, single iPSCs^7^ or neural precursor cells (NPCs)^9,10^ into synthetic or cell-derived matrices, such as Matrigel. However, the brain extracellular matrix (ECM) has a unique composition and properties that cannot yet be reproduced synthetically. As the brain ECM influences cell functionality, maturation^11^, and likely also pathogenesis, reproduction of a physiological environment may be required to establish brain-like cell states, functions, and disease responses, thus limiting the applicability of hydrogel-based systems. Spheroid cultures are commonly derived by aggregation of NPCs^12^ or NPCs and astrocytes^13^, followed by further maturation of the cells in 3D. Although these models show good reproducibility when generated in microwells, they lack stable incorporation of mature and homeostatic microglia (MG), the brain-resident immune cells, and long-term *in vitro* maintenance beyond a few weeks. Thereby, recapitulation of tissue characteristics is limited, including formation and maturation of ECM, and establishment of mature morphologies, functions, and interactions of the different cell types. In contrast, NO develop in 3D directly from stem cells and can thus uniquely model aspects of human brain development^14,15^, thereby giving important insights into human-specific mechanisms, e.g., regarding precursor cell biology and cell-type specifications^16^. However, NO cultured *in vitro* show limited suitability to study MG, which play major roles in maintenance of brain tissue physiology and neurodegenerative diseases^17^. MG arise from the mesoderm and are therefore not observed in neuroectodermal organoids derived with guided protocols, whereas unguided protocols can yield few MG if mesodermal cells are present^18^. Exogenously added iPSC-derived microglia (iMG), although promoting organoid maturation^19^, rapidly decline in numbers and usually disappear within few weeks^20,21^. Furthermore, these iMG do not show mature and homeostatic signatures, including cell surface expression of P2RY12 protein, or morphologies *in vitro*^19,20,22–24^. Such physiological features can however be achieved by xenotransplantation of NO with iMG into mouse brain^20^. Even though these models are highly complex and low-throughput, limiting their suitability for drug testing, they confirm the fundamental ability of iMG to mature in environments provided by iPSC-derived models.

We therefore hypothesized that crucial extrinsic factors are required for induction of a mature, homeostatic MG state, which are not found in currently available human 2D and 3D *in vitro* models. These factors may include a post-mitotic, mature brain-parenchyma-like environment, low levels of cell death and associated stress, and reproducible and continuous presence of mature astrocytes. The inapt environment provided by current models, such as the lack of mature astrocytes in trackable time frames^25,26^ and the proliferative state with neural rosettes, associated large-scale growth and subsequent necrotic core formation^27,28^ present in NO cultured *in vitro*, may thus prevent formation of a mature iMG phenotype. Furthermore, the artificial, environment-related activation observed in non-homeostatic iMG *in vitro* may impair studies on microglial responses to brain disease, such as AD.

To overcome drawbacks of current models, we developed a highly reproducible, iPSC-derived 3D cortical tissue model that shows high viability for many months in a postmitotic state without necrotic core formation, with efficient and reproducible incorporation of cortical neurons (iNE), astrocytes (iAS), and iMG. We show that co-cultured cells form a 3D structure with several tissue characteristics, including formation and maturation of a brain-like ECM, as well as maturation of cell types on a morphological, functional, and protein expression level, and adoption of a mature iMG state with homeostatic marker expression and ramified morphology. Using CRISPR/Cas9 genome editing, we established a human AD model based on knock-in of synergistic APP mutations, which displayed disease-associated features including amyloid deposition, increased phospho-tau levels, neuroinflammation and altered microglial expression signatures relevant for AD. We further show formation of Aβ plaque-like structures upon addition of exogenous Aβ and confirm suitability of the model to test anti-amyloid antibodies. Taken together, we developed the – to our knowledge – first fully iPSC-based 3D *in vitro* model of brain tissue that allows studying physiological and pathological phenotypes of mature neurons, astrocytes, and especially microglia in a reproducible, human tissue-like environment.

## Results

### Generation of a reproducible human 3D brain tissue model containing homeostatic microglia

We hypothesized that generation of a reproducible and controllable 3D cortical brain tissue model (3BTM) can be achieved by combining and aggregating brain cell precursors that have achieved a stable fate but are still plastic enough to allow for tissue integration (**Figure 1A**). Starting from these differentiated cells should reduce undesired heterogeneity compared to systems originating from stem cells or early progenitor cell populations. To test this approach, we first optimized attached, small-molecule-based, 2D differentiation protocols to separately obtain iNE^29^, iAS^30^ and iMG^31^ from human iPSCs (**Extended Data Fig. 1A**), to promote homogeneous differentiation and culture purity. We confirmed typical morphologies, marker expression, and purity of the different cell types by immunostainings (**Extended Data Fig. 1B-D**). Next, the differentiated cells were self-aggregated into 3D assemblies by plating into ultra-low attachment plates (**Figure 1A**). We avoided the use of exogenous ECMs, such as Matrigel, to minimize variability of exogenous cues, and thus increase reproducibility and simplicity. We combined iNE and iAS in a 3:1 ratio, similar to human cortex^32^, and confirmed cell aggregation into uniform, sphere-like structures of about 800 µM diameter (**Figure 1B, 1C**). To avoid culture growth and necrotic core formation and reach a stable, postmitotic state rapidly after cell aggregation, cultures were treated with DAPT, a Notch-signaling inhibitor that drives residual NPCs into terminal neuron differentiation^33^, followed by 5-Fluorouracil (5FU), a pyrimidine analog that kills residual dividing cells^29^ (**Figure 1A**). The combination of optimized cell differentiations, time points for cell aggregation, and compound treatments yielded 3BTMs with largely stable diameters over several months in culture without relevant growth or shrinkage (**Figure 1C**, 95% confidence intervals: 1 month: 779-802 µm, 3 months: 771-788 µm, 6 months: 766-783 µm). Further confirming efficacy of the applied compound treatments in inducing a post-mitotic state, we found that proliferating cells were abundant on Day 3 after culture generation, before 5FU treatment, but almost completely absent at all later timepoints tested (**Figure 1D**). In addition, cultures maintained the initial iNE:iAS ratio of 3:1 within and across independent experiments over time (**Figure 1E**), again demonstrating reproducibility and stability of the selected cell ratios. Both iNE and iAS formed dense and uniformly distributed networks as seen by immunostainings for marker proteins (**Figure 1F**), indicating mature cell morphologies and absence of necrotic core formation. Hypoxia, which usually contributes to necrosis, was largely absent in 3BTMs grown under normoxic conditions, in contrast to 3BTMs cultured under hypoxic conditions (**Extended Data Fig. 1E**). To increase cellular complexity and physiological relevance of the cultures and enable analysis of microglial states and behavior, we next added iMG to 3BTMs (**Figure 1G**). Added iMG migrated into the 3D cultures autonomously within a few days (**Extended Data Fig. 1F**) and spread within the cultures (**Figure 1H, Extended Data Fig. 1G**), resembling the tissue tiling seen in brain. iMG incorporation was strongly improved in the presence vs. absence of iAS (**Extended Data Fig. 1H)**. Triple co-cultures could be maintained for at least 6 months without necrotic core formation, and with iNE and iAS maintaining densely branched networks (**Extended Data Fig. 1I**). Strikingly, iMG adopted an increasingly ramified morphology (**Figure 1I, 1K**) and expressed the homeostatic marker P2RY12 in the membrane of over 75% of cells at 6 months, suggesting a broadly present homeostatic state *in vitro* (**Figure 1L**). Importantly, such homeostatic microglial phenotypes have not been observed in organoids cultured *in vitro* but were only achieved by transplantation *in vivo*^20^. iMG were present in 3BTMs for over 6 months and slowly decreased over time (**Extended Data Fig. 1K**), possibly due to senescence^34^ or migration out of the culture towards a growth factor gradient into the media. However, survival was much higher than in current *in vitro* models^20,21^, enabling analysis of physiological and neuroinflammatory phenotypes at advanced ages of 3 to 6 months. In addition, we found that iMG can be incorporated into 3BTMs at later timepoints, e.g., at 5 months, leading to a dense, ramified population at 6 months of age (**Extended Data Fig. 1L, M**). P2RY12 expression was similar to iMG at 1 month (**Extended Data Fig. 1N**). Next, we analyzed cell death in 3BTMs using TUNEL stainings, and found consistently low levels, also in long-term cultures, especially when all three cell types were present (**Figure 1M, Extended Data Fig. 1O**), suggesting that iMG not only benefit from but also contribute to the maintenance of a physiological tissue environment, e.g., by phagocytosis of potentially toxic degradation products, dead cells, or cell debris. Finally, we used an independent iPSC line to confirm reproducibility of 3BTM generation across cell lines. We repeated analyses of size (**Extended Data Fig. 1P**), iNE:iAS ratio (**Extended Data Fig. 1Q**), post-mitotic state (**Extended Data Fig. 1R**) and cell morphology (**Extended Data Fig. 1S**), which produced highly similar results. In summary, we established a highly reproducible, controllable, mature, and stable 3D cortical tissue model with efficient and long-lasting incorporation of iNE, iAS and iMG.

**Figure 1.**
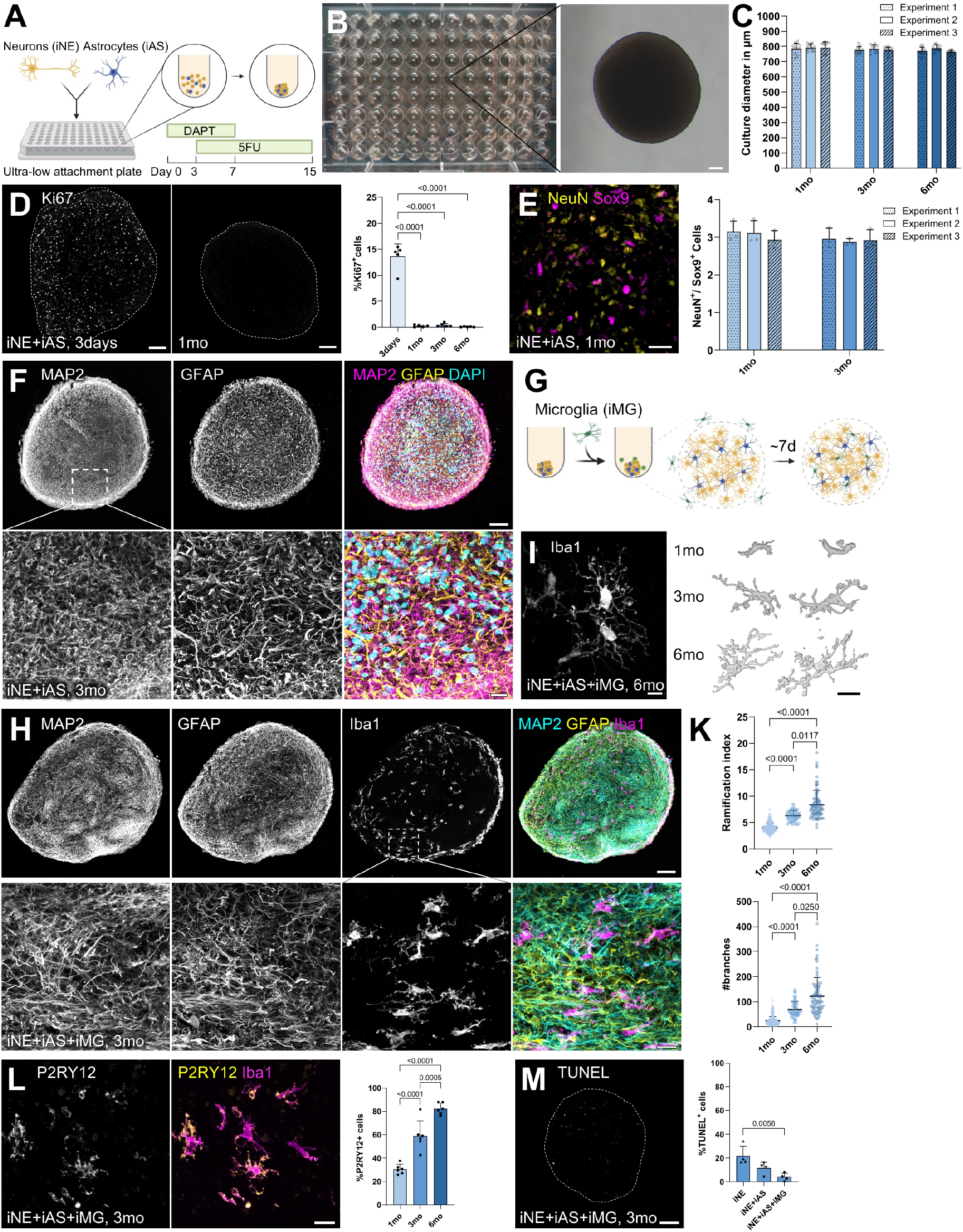
Generation of a reproducible human 3D brain tissue model containing homeostatic microglia. **(A)** Experimental approach to generate 3BTMs by self-aggregation of iNE and iAS in ultra-low attachment plates. **(B)** Overview of a 96-well plate with 3BTMs at 1 month (1mo) of age, note uniform size and shape of the cultures. Right: Brightfield image of a 3D culture at 1 month, scale bar 100 µm. **(C)** Quantification of culture diameters at 1, 3 and 6 months (mo) (n = 3 experiments with 10-11 3BTMs per experiment and timepoint, each dot represents one 3BTM). **(D)** Immunofluorescence (IF) stainings (left) and quantification (right) of Ki67-positive cells in 3D co-cultures over time (n = 5 experiments with one 3BTM each, each dot represents one experiment, one-way ANOVA with Tukey’s multiple comparisons test), scale bars 100 µm. **(E)** IF staining showing NeuN-positive iNE and Sox9-positive iAS in 3D co-cultures (scale bar 30 µm). Quantification of the NeuN:Sox9 ratio at 1 and 3 months on the right (n = 3 experiments with three 3BTMs per experiment and timepoint, each dot represents one 3BTM). **(F)** IF staining of a middle slice from an iNE+iAS culture at 3 months for MAP2 (iNE) and GFAP (iAS), scale bar 100 µm. Zoom-in at the bottom (scale bar 20 µm). **(G)** Experimental approach to generate 3BTMs containing iNE, iAS, and iMG. iNE and iAS are aggregated as in (A). iMG are added 15 days later into the media and migrate into the cultures over ∼7days. **(H)** IF staining of a middle slice from a 3-month-old iNE+iAS+iMG culture showing neuronal (MAP2) and astrocytic (GFAP) networks and distribution of incorporated iMG (Iba1), scale bar 100 µm. Zoom-in showing cell morphologies with fine and dense processes of iNE and iAS, and ramifications of iMG, scale bar 20 µm. **(I)** Left: IF staining of iMG at 6 months showing highly ramified morphology, scale bar 10 µm. Right: Representative 3D reconstruction of MG in 3BTMs at 1/3/6 months of age scale bar 10 µm. **(K)** Quantification of MG morphology measured by ramification index (top) and number of branches (bottom) per cell at 3 timepoints (n = 3 experiments with 14-60 cells from 2-3 3BTMs per experiment and timepoint, each dot represents one iMG, Kruskal-Wallis test with Dunn’s multiple comparisons test). **(L)** IF staining at 3 months for homeostatic membrane marker P2RY12, scale bar 20 µm. Quantification of P2RY12-positive cells over time on the right (n = 5-6 experiments with 26-159 cells from two 3BTMs per experiment and timepoint, each dot represents one experiment, one-way ANOVA with Tukey’s multiple comparisons test). **(M)** IF staining showing levels of cell death by TUNEL staining in iNE+iAS+iMG cultures at 3 months, scale bar 100 µm. Quantification on the right showing comparison of different cell type combinations (n = 4 experiments with two 3BTMs per cell-type combination, each dot represents one experiment, one-way ANOVA with Tukey’s multiple comparisons test). Figures 1A and 1G were created in BioRender. Data are presented as mean + SD.

### Maturation of 3BTMs increases similarity to human brain tissue

Our data suggested that 3BTMs provide a stable and postmitotic environment promoting iMG maturation over time. Thus, we hypothesized that all incorporated cell types may develop similar phenotypes to their counterparts *in vivo*, resulting in a high degree of maturation. To investigate this, we first performed mass spectrometry analysis of 3BTM proteomes at 3 days, 3 months, and 6 months after culture generation, and compared them to neural organoid (Day 90), fetal human brain (FB, gestational weeks (GW) 15-17) and juvenile human brain (JB, 6 yrs) samples. Principal component (PC) analysis (**Figure 2A, Extended Data Fig. 2A**) resulted in a dense clustering of samples of similar ages and replicates, showing high reproducibility of the cultures regarding overall protein composition. This was confirmed by strong correlation between sample replicates (**Extended Data Fig. 2B**), with biological replicates of 3BTMs showing coefficients highly similar to technical replicates of the neural organoid and juvenile brain samples. We next focused on characterizing PC1, explaining most of the observed variance across tested models. Interestingly, samples separated along PC1 in a putative maturation-dependent manner, with neural organoids and 3-day-old samples on one end, and 3- and 6-months-old 3BTM samples placed between the fetal and juvenile human brain samples (**Figure 2A)**. Gene ontology enrichment analysis for biological processes (GO-BP) indeed confirmed that upregulated proteins were involved in maturation-dependent, synapse-associated biological processes such as chemical synaptic transmission and ion/neurotransmitter transport, whereas downregulated proteins were involved in cell proliferation and DNA replication-related processes (**Figure 2B**). We focused on PC1, as PC2 explained much less of the variance in the dataset (**Extended Data Fig. 2A**) and was largely driven by “blood-dependent” processes or DNA replication not relevant for matured 3BTMs (**Extended Data Fig. 2C**).

**Figure 2.**
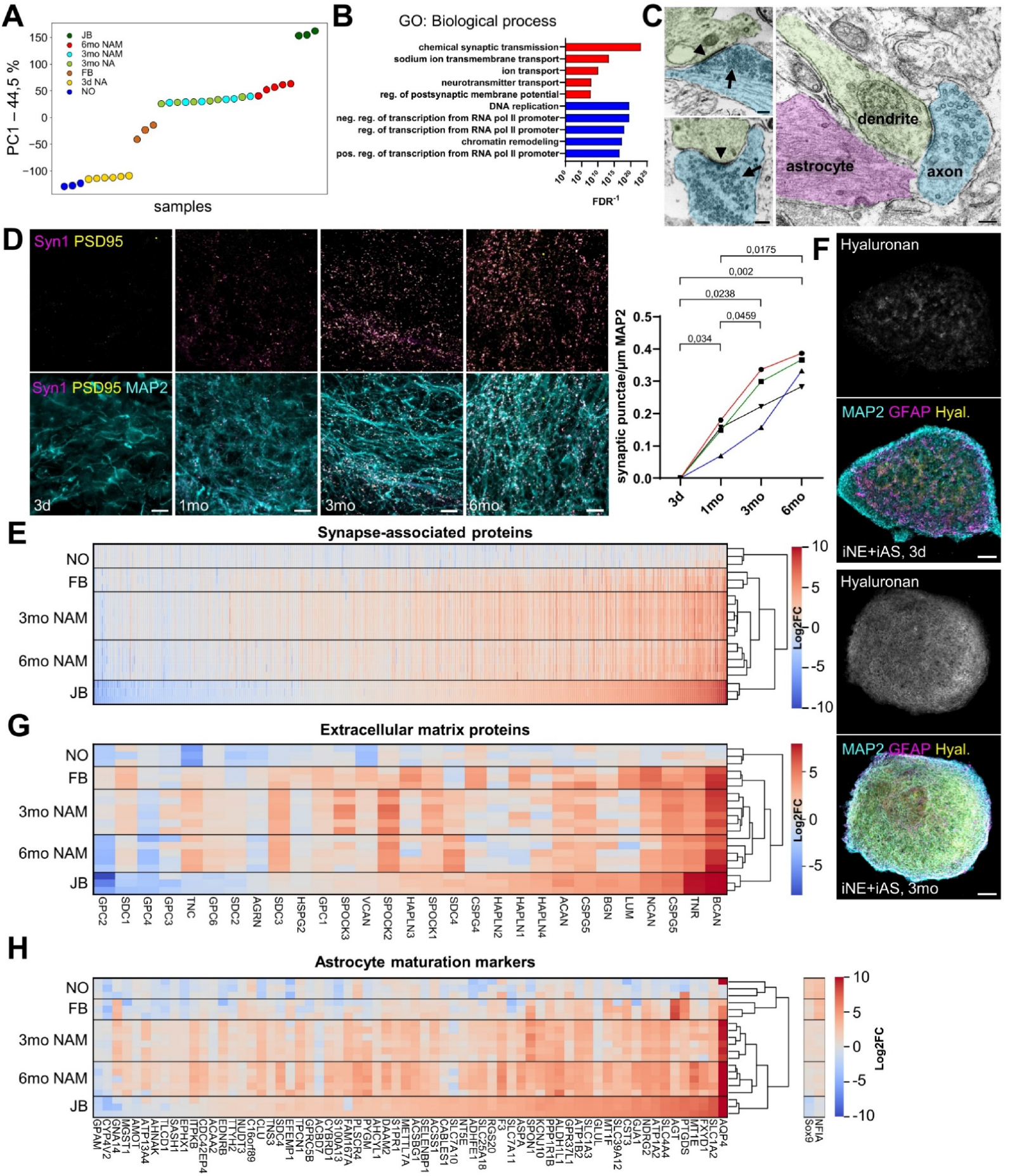
Maturation of 3BTMs increases similarity to human brain tissue. **(A)** Principal component analysis (PCA) of proteomic dataset based on all proteins after imputation. Shown is principal component 1 (PC1), explaining 44.5% of the variability in the dataset. NO = neural organoid sample (Day 90, 3 technical replicates), 3d NA = 3-day-old 3BTMs of iNE+iAS (6 biological replicates), FB = fetal brain (GW 15-17, 3 biological replicates), 3mo NA = 3-months-old 3BTMs of iNE+iAS (6 biological replicates), 3mo NAM = 3-months-old 3BTMs of iNE+iAS+iMG (6 biological replicates), 6mo NAM = 6-months-old 3BTMs of iNE+iAS+iMG (5 biological replicates), JB = juvenile brain (6 yrs, 3 technical replicates). **(B)** Gene ontology (GO-BP) analysis of top proteins influencing PC1, shown are processes in which upregulated proteins (red) or downregulated proteins (blue) are enriched, sorted by false-discovery rate (FDR). **(C)** Electron micrographs of an iNE+iAS+iMG culture at 3 months showing formation of synapses (left, scale bars 200 nm) with presynaptic terminals (arrows, pseudocolored in blue) and postsynaptic densities (arrowheads, pseudocolored in green), and a tripartite synapse (right, scale bar 200 nm) made of presynaptic axon, postsynaptic dendrite, and adjacent iAS (pseudocolored in purple). **(D)** IF stainings of iNE+iAS+iMG cultures over time for presynaptic Synapsin-1 (Syn1) and postsynaptic PSD-95, scale bars 10 µm. Quantification of colocalized puncta, suggesting synapse formation, normalized to MAP2 over time (n = 4 experiments with two 3BTMs each represented by colored lines, repeated-measures one-way ANOVA with Tukey’s multiple comparisons test). **(E)** Heatmap showing log_2_-fold changes (log_2_FC) in protein levels of synapse-associated proteins found in proteomics datasets compared to 3-day-old iNE+iAS cultures (set to zero). Hierarchical clustering of the different samples shown on the right. **(F)** IF stainings of iNE+iAS cultures at 3 days and 3 months showing deposition of Hyaluronan in 3BTMs, scale bars 100 µm. **(G)** Heatmap and clustering as in (E) showing ECM proteins. **(H)** Heatmap and clustering as in (E) showing astrocyte maturation markers and housekeeping/identity proteins Sox9 and NFIA.

To confirm relevance of the identified pathways and maturation of 3BTMs over time, we analyzed synapse formation as the main process driving PC1. We performed electron microscopy analysis and indeed found differentiated synapses (**Figure 2C, left**), and in some cases also brain-typical tripartite synapses (**Figure 2C, right**), suggesting not only connections between iNE but also neuron-glia interactions. In addition, immunostainings for presynaptic Synapsin-1 and postsynaptic PSD95 revealed increased co-localization over time (**Figure 2D**), suggesting synapse formation. To get a more global overview of the synaptic machinery, we analyzed synapse-associated proteins in our proteomic dataset and found strong upregulation of most proteins in 3- and 6-month-old cultures (**Figure 2E**), positioning them closer to fetal and juvenile brain samples than to organoids or the 3-day-old cultures used for comparison. We next confirmed downregulation of cell-cycle-associated proteins in older cultures, again indicating higher maturity and similarity to the juvenile brain sample and confirming the post-mitotic state of the cultures (**Extended Data Fig. 2D**). The organoid sample did not differ considerably from the 3-day-old 3BTMs used for comparison, indicating a lower maturation level. Interestingly, “cell adhesion” was another significantly altered process in our GO-BP analysis for PC1, and “extracellular region” and “extracellular space” were enriched processes in a GO analysis for cellular compartments. Therefore, and to further investigate maturation markers indicating tissue-related properties, we analyzed formation of a brain-like ECM in 3BTMs. The brain ECM is essential for a variety of cell functions, such as network formation and signaling events^11^, however, due to its distinct properties, it so far could not be replicated *in vitro*^35^. We therefore refrained from adding any exogenous ECM when generating the cultures and rather hypothesized that the cells would produce their own, brain-like matrix. Strikingly, we found that hyaluronan, a main component of brain ECM, was first present at low, disperse levels on day 3 and strongly increased at 3 months to a homogeneous staining (**Figure 2F**) sparing cell bodies (**Extended Data Fig. 2E**), indicating deposition in an ECM-like structure. Hyaluronan levels were strongly increased in the presence of iAS (**Extended Data Fig. 2F**), suggesting that iAS are the main producers. To get a global overview of ECM formation, we analyzed ECM-related proteins in our proteomics dataset. We detected a variety of ECM molecules, including brain-specific or -enriched core proteins of chondroitin sulphate proteoglycans, such as Neurocan (NCAN), Brevican (BCAN), and CSPG5, linker proteins Tenascin (TNC) and HAPLNs, and other relevant molecules such as hyaluronan sulphate proteoglycans (**Figure 2G**). We found a strong upregulation of many brain ECM proteins in our 3-month-old and 6-month-old samples, again making them more similar to the fetal and juvenile brain samples, compared to the 3-day-old and neural organoid samples (**Figure 2G**). Moreover, we found a set of downregulated proteins in aged cultures that matched reduced expression in juvenile vs. fetal brain, e.g., Glypican-2 (GPC-2) and -4 (GPC-4), especially at 6 months, suggesting not only build-up but also brain-like remodeling of the cultures’ ECM over time.

As astrocytes are a main driver of brain tissue maturation^36^, we analyzed our proteomics dataset to determine levels of mature human astrocyte markers described previously^37^. We found a strong upregulation of most astrocyte-specific maturation markers in 3- and 6-month-old cultures, including Aldh1l1, Aqp4, and excitatory amino acid transporters, whereas housekeeping proteins NFIA and Sox9 stayed largely constant over time (**Figure 2H**). This led to a higher similarity to the juvenile than to the fetal brain sample as seen by hierarchical clustering, showing the high maturity of iAS in 3BTMs. To further study iAS maturity, we incorporated a small percentage of membrane-eYFP-expressing iAS into the cultures to investigate single cell morphologies. We found several different types of iAS with intricate morphologies, including highly ramified, “bushy” cells resembling protoplasmic astrocytes in human brain^38^, less ramified cells extending several processes over 200-300 µm distance, and very large cells extending few processes over the entire culture diameter (**Extended Data Fig. 2G**). Similarly, we investigated single iNE morphology after transduction with AAVs expressing eGFP under a human Synapsin-1 promoter. Again, we found a mature cell morphology with a branched dendritic tree and long, ramified axon (**Extended Data Fig. 2H**), resembling pyramidal neurons in the brain. Together, our analysis confirms that iNE and iAS in 3BTMs acquire mature morphologies, protein composition, and ECM formation, similar to juvenile brain cells *in vivo*.

### iMG mature in 3BTMs and adopt transcriptomic signatures similar to human brain

Given that the ramified iMG morphology and expression of markers like P2RY12 suggested iMG maturation and homeostasis (**Figure 1K**), we performed single-cell RNA sequencing analysis (scRNAseq) to assess iMG gene expression profiles after prolonged integration into 3BTMs (**Figure 3A**). To study phenotype maturation over time, we analyzed cells from two different experiments at 1 and 3 months of age. As we were most interested in iMG states, we chose a gentle dissociation protocol including Actinomycin D as a transcriptional inhibitor to promote retrieval of the cells without artificial activation^39^. Consequently, after quality filtering (**Extended Data Fig. 3A**), the iMG-optimized extraction resulted in successful recovery of iMG, but only very few iNE and iAS (**Extended Data Fig. 3B**), which due to their highly complex structure and strongly interconnected networks likely require more harsh dissociation procedures for efficient retrieval. We excluded iNE, iAS, and mixed clusters from the subsequent analysis, and focused on the wildtype (WT) iMG subset at 1 and 3 months (**Extended Data Fig. 3C**). After filtering for high-quality cells (**Extended Data Fig. 3D**) we identified a total of 8,827 iMG mainly separating by the age of the source cultures. Cluster composition and UMAP from the two independent experiments were highly similar, demonstrating reproducibility of iMG states in 3BTMs (**Extended Data Fig. 3E**). Furthermore, upon clustering of the UMAP space, similar major cell states were identified at 1 and 3 months. To annotate these cell states, we employed cluster marker genes (**Extended Data Fig. 3F**) and their functional enrichment within literature-described^40^ existing phenotypes *in vitro* and in humans. We thereby identified *resting* and *chemokine-associated* clusters (based on marker gene expression (**Extended Data Fig. 3F**) present at both 1 and 3 months and additional *proliferative* and a small *glycolytic* cluster present at 1 month (**Figure 3B, 3C**). Across all clusters, typical MG markers such as AIF1, TYROBP, and HEXB were expressed (**Figure 3D**). Differential gene expression analysis (DEA) yielded 779 up- and 618 downregulated genes in 3 months compared to 1 month *resting* iMG, suggesting a phenotypic transition with culture age as hypothesized (**Figure 3E, F**). Gene set enrichment analysis (GSEA) of DEGs pointed towards a metabolic shift over time, as well as an upregulation of genes involved in immune-related functions such as “MHC class II protein complex assembly”, “receptor-mediated endocytosis” and “antigen processing and presentation” (**Figure 3G**). These findings suggest the acquisition of an immune-sensing state over time, a hallmark of MG maturation enabling the cells to surveil the environment and react to aversive cues^41^. We therefore compared a previously published sensome signature^42^ to our iMG signatures at 1 and 3 months and found that iMG at 3 months had a significantly higher enrichment for the sensome signature, indicating increased maturity and ability to interact with their environment (**Figure 3H**). Pseudotime analysis further validated the phenotypic progression of iMG from 1 to 3 months in culture, as highlighted by the inferred trajectory (**Figure 3I**). Starting from *proliferative* cells at 1 month, the trajectory extended towards the intermediate 1 month *resting* state and further towards the 3 months states, as expected. Further supporting the cell states’ annotation suggested in Figure 3B for 1-month iMG, pseudotime analysis of the 1mo subset highlighted an unfolding of phenotypes starting from a *proliferative* state, passing through the *resting* state, and then branching into the *activated* and *glycolytic* clusters. (**Figure 3I**). To obtain more detailed information on iMG maturation and compare the cells to MG from human brain, we next integrated our dataset with a published dataset of postmortem human fetal (GW 10-14) and postnatal MG (14-23 yrs)^43^ and assessed individual cells’ enrichment for the adult MG signature published by Galatro et al.^44^. Intriguingly, iMG from 3-months-old cultures clustered along postnatal MG, and iMG from 1 month to 3 months showed a similar transition towards a more mature phenotype as observed from fetal to postnatal MG, attested by the increased enrichment for the Galatro signature with age in both datasets (**Figure 3K**). As these insights suggest the progressive acquisition of a postnatal rather than fetal profile at 3 months in culture^41^ we next compared our iMG to fetal and adult human MG signatures from an independent study^45^, after confirming the suitability of the approach with another post-mortem dataset^46^ (**Extended Data Fig. 3G**). We found a strongly increased enrichment of our 3-month iMG for the adult compared to fetal signature (**Figure 3L)**, again demonstrating the maturity of iMG in 3BTMs. Finally, we compared our iMG signature at 3 months to different human *in vitro* model signatures derived from a published dataset^21^ (**Extended Data Fig. 3H)**, comprising primary fetal MG cultured in 2D or incorporated into neural organoids, and iMG cultured in 2D or xenotransplanted into mouse brain. We found the strongest enrichment for xenotransplanted iMG (**Figure 3M**), which more closely resemble *in vivo* MG^47–49^. Taken together, our scRNAseq analysis strongly supports that the tissue-like environment in 3BTMs induces maturation of iMG globally on transcriptomic level, making them increasingly similar to their *in vivo* counterparts over time.

**Figure 3.**
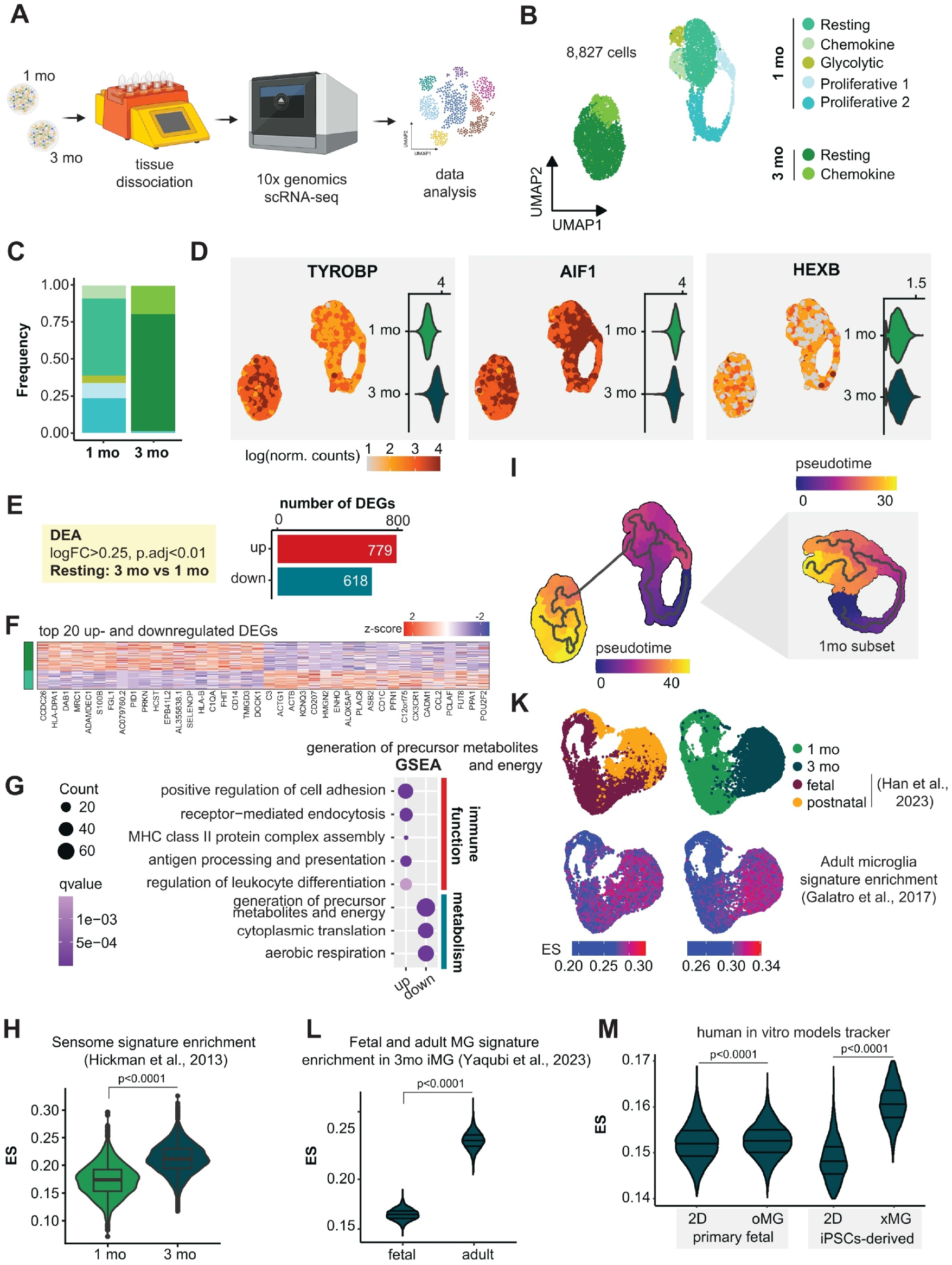
iMG mature in 3BTMs and adopt transcriptomic signatures similar to human brain. **(A)** Schematic overview of the single-cell RNA-sequencing experiment for the investigation of iMG phenotypes in 3BTMs at 1 and 3 months. iMG from two independent experiments were used. Created in BioRender. **(B)** UMAP representation (res=0.3, dims = 1:10) of 8,827 cells and cluster annotation of iMG at 1 and 3 months of age. **(C)** Cell state frequencies reported for iMG at 1 and 3 months separately. **(D)** Normalized expression of key MG markers (*TYROBP, AIF1, HEXB*) both UMAP-projected and as a violin plot in iMG at 1 and 3 months. **(E**) Number of differentially expressed (DE) genes (p.adj<0.01, abs(log_2_FC)>0.25) obtained by comparing 1 month and 3 months *resting* clusters. **(F)** Heatmap showing normalized scaled expression of the top 20 up- and downregulated DEGs. **(G)** Over-representation analysis (ORA, gene ontology, biological process) of DEGs showing enrichment of upregulated genes in immune-function-related processes and of downregulated genes in metabolism-related processes (*q*-value<0.2). **(H)** Violin plot showing MG sensome signature^42^ enrichment scores (ES) for iMG at 1 and 3 months (Wilcoxon signed-rank test). **(I)** Pseudotime analysis highlighting phenotypic progression of iMG from 1 month to 3 months. 1 month subset on the right. **(K)** Top: UMAP plot after integration of a human fetal and postnatal, *ex vivo* MG dataset (Han et al.^43^) with iMG at 1 and 3 months, split by dataset. Bottom: Same UMAP plot as above showing projected adult MG signature (Galatro et al.^44^) ES, split by dataset. **(L)** Violin plot reporting fetal vs. adult MG signature (extracted from Yaqubi et al.^45^) ES in iMG at 3 months. **(M)** *In vitro* tracker reporting ES of iMG at 3 months across different model-specific signatures derived from the integrated dataset by Popova et al.^21^, including primary human fetal MG cultured in 2D (2D, primary fetal) or incorporated into cortical organoids (oMG, primary fetal) as well as human iMG cultured in 2D (2D, iPSCs-derived) or xenotransplanted into mouse brain (xMG, iPSCs-derived) (Wilcoxon signed-rank test). Data in box plots are presented as median, 25^th^ and 75^th^ percentiles (box) ± 1.5x IQR (whiskers).

### iNE, iAS, and iMG in 3BTMs acquire functionalities typical for mature brain cells

We next investigated if the observed adoption of more mature states on multiple levels would also lead to a functional maturation of the cells. As we observed extensive synapse formation and expression of various synapse-related proteins and ion channels, we first performed electrophysiological patch-clamp measurements to analyze iNE functionality. We observed spontaneous action potentials (**Figure 4A, Extended Data Fig. 4A**) and could evoke action potentials in iNE by current injection (**Figure 4B**), which was blocked by application of tetrodotoxin (**Figure 4C**), demonstrating dependence on voltage-gated sodium channels. We further observed heterogeneous firing patterns in response to depolarizing stimuli in different iNE (**Extended Data Fig. 4B**), possibly indicating presence of various neuronal subtypes as described for the human cortex^50^. Finally, cells showed a resting membrane potential of around -65 mV (**Figure 4D**), as shown before for mature neurons in human cortex^50^. Altogether, the data suggest that 3BTMs contain mature iNE with synapses that express typical ion channels underlying intrinsic active and passive neuronal membrane properties. To analyze functionality and network formation on a larger scale, we performed whole-mount calcium imaging using Fluo-4 and detected spontaneous and synchronized calcium waves (**Extended Data Fig. 4C, Supplementary Video 1**), suggesting functional network formation.

**Figure 4.**
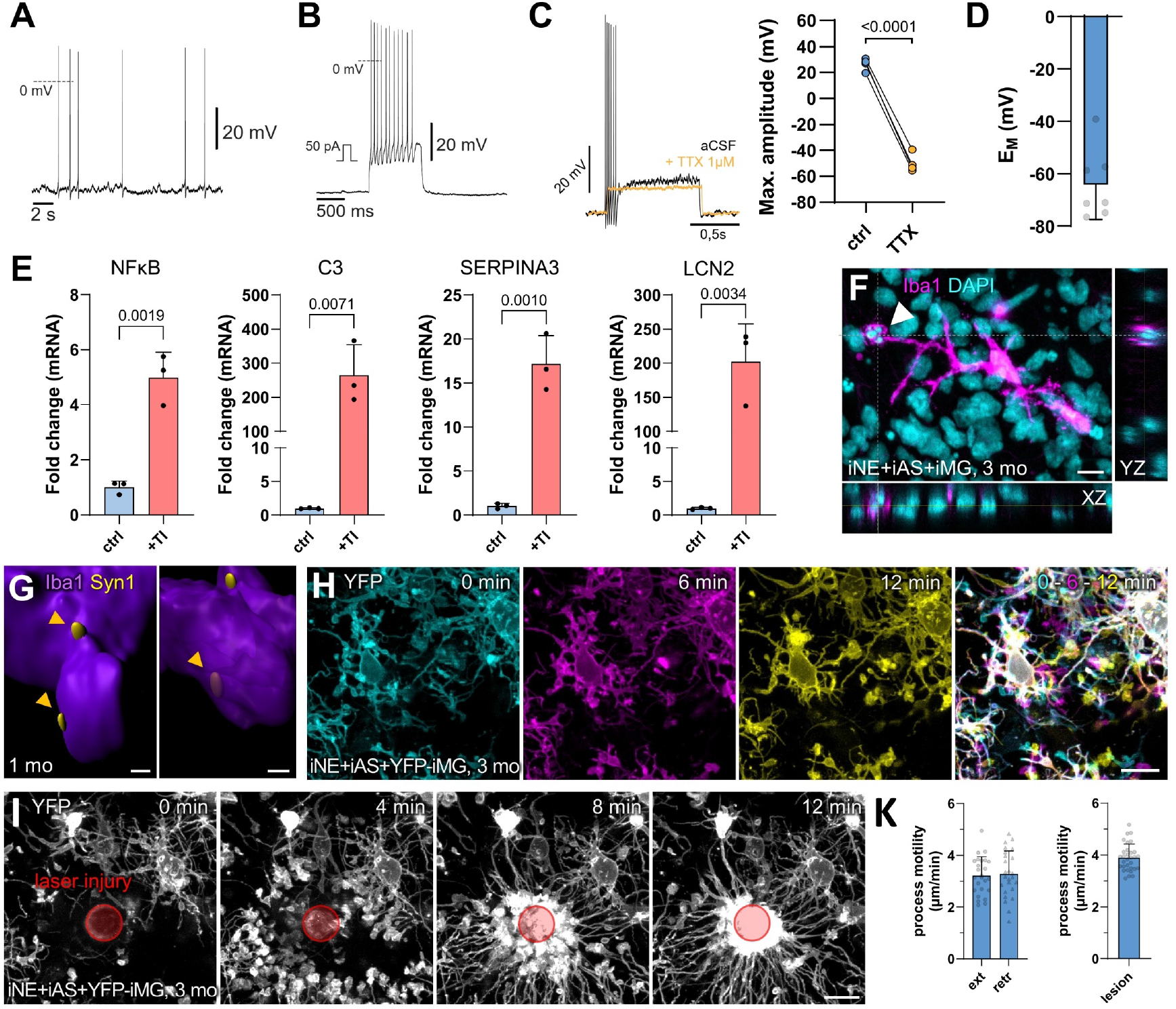
iNE, iAS, and iMG in 3BTMs acquire functionalities typical for mature brain cells. **(A)** Representative patch-clamp measurement of single iNE in iNE+iAS+iMG 3BTMs at 3 months showing spontaneous action potentials. **(B)** Representative patch clamp measurements as in (A) with current injection showing evoked action potentials. **(C)** Application of tetrodotoxin (TTX) inhibits action potential firing, quantification on the right (n = 2 experiments with 2 cells per experiment, each dot represents one cell, paired two-sided *t*-test). **(D)** Resting membrane potential (E_M_) measurements of single iNE as in (A) (n = 2 experiments with 3-4 cells per experiment, each dot represents one cell). **(E)** qPCR analysis for astrocytic activation markers in iNE+iAS cultures at 1 month of age treated with PBS (ctrl) or TNFα+IL1α (TI), normalized to *SOX9* as an astrocytic reference gene (n = 3 experiments with 7-10 3BTMs pooled for RNA extraction, each dot represents one experiment, unpaired two-sided t-tests). **(F)** IF staining for DAPI-positive DNA and Iba1-positive iMG in 3-month-old 3BTM showing microglial process engulfing a dead cell’s nucleus (as seen by bright and fragmented DAPI staining, arrowhead), orthogonal projections in YZ and XZ, scale bar 10 µm. **(G)** 3D surface rendering of IF staining for Synapsin-1 (Syn1)-positive synaptic material and Iba1-positive iMG at 1 month showing close association (left) and engulfment (right) of synaptic material by iMG, scale bars 0,5 µm. **(H)** 2-Photon live-cell imaging of iNE+iAS+iMG 3BTMs at 3 months with iMG expressing membrane-eYFP. Pictures showing different, color-coded timepoints (0, 6, 12 min), merged image on the right showing movement by color-coding whereas stable structures appear white, scale bar 20 µm. **(I)** 2-Photon live-cell imaging as in (H) showing reaction of iMG to a focal laser injury (red circle), scale bar 20 µm. **(K)** Quantification of iMG process motility under baseline conditions (left, ext = extension, retr = retraction, n = 3 experiments with 8-9 tracks per experiment and direction, each dot represents one track) and after focal laser injury (right, n = 3 experiments with 10-11 tracks per experiment, each dot represents one track). Data in bar graphs presented as mean + SD.

To investigate iAS functionality, we focused on the cells’ ability to react to an activating stimulus, such as inflammatory cytokines. We treated 3BTMs made of iNE and iAS with TNFα and IL1α as described previously^51^ to promote a reactive astrocyte state. After 24 hours, we found a robust increase in typical inflammation and reactive astrocyte markers *NFκB, C3, SERPIN3A*, and *LCN2* (**Figure 4E**) by qPCR, confirming the cells’ ability to switch to an activated state.

Finally, we determined functionality of iMG. As innate immune cells, iMG constantly surveil the environment with their processes, remove debris and excess synapses, and immediately react to adverse stimuli such as tissue injury^52^. To investigate these functions, we first performed immunostainings for iMG and either cell nuclei or pre-synaptic Synapsin-1 and indeed observed iMG engulfing a nucleus of a dead cell, as seen by bright and fragmented DAPI staining (**Figure 4F**), as well as engulfing or closely contacting synaptic material (**Figure 4G, Supplementary Video 2**). We next analyzed membrane-eYFP-expressing iMG in 3BTMs at 3 months using 2-photon live-cell imaging. At baseline conditions, iMG constantly surveilled their environment using their ramified processes (**Figure 4H, Supplementary Video 3**). Strikingly, when we inflicted a focal laser injury into the culture, iMG processes from all surrounding cells immediately extended towards the lesion, leading to a complete ensheathment of the lesion site after around 10-12 minutes (**Figure 4I, Supplementary Video 4**), whereas the cell bodies remained in their original position. When quantifying movement speed of cell processes, we found similar values for extension and retraction under baseline conditions, and higher speeds after the injury (**Figure 4K**), consistent with previous *in vivo* data^20,41^. This shows that the iMG can launch a classical injury response similar to cells *in vivo* and confirms presence of functional P2RY12 on the cell surface, which is required for this rapid response^53,54^. Altogether, all cell types gained brain-like functionalities indicating functional maturation in 3BTMs.

### CRISPR/Cas9-mediated insertion of synergistic APP mutations to model Alzheimer’s disease

Since our model recapitulated several morphological and functional aspects of mature brain tissue, we next aimed to test if it can be applied to induce hallmarks and study mechanisms of Alzheimer’s disease, such as accumulation of Aβ and neuroinflammation. We therefore knocked in a combination of synergistic, AD-causing mutations into WT iPSC using CRISPR/Cas9 gene editing to induce rapid pathogenesis while avoiding overexpression artefacts. This strategy has already been successfully applied in mouse models of AD, and the resulting NLGF knock-in (KI) mice^55^ are broadly used in the field. To replicate the NLGF model, we started from a previously published line carrying the APP Swedish (NL) mutations^56^ and added APP Arctic (G) and Iberian (F) mutations (**Extended Data Fig. 5A**), which accelerate protofibril formation and increase Aβ42:40 ratio, respectively. The resulting NLGF-KI iPSC line was quality controlled for successful editing of all mutations by Sanger sequencing (**Extended Data Fig. 5B**), presence of both alleles of the edited locus^57^ and absence of trisomy at the *BCL2L1* locus (**Extended Data Fig. 5C**), a common pro-survival aberration found in cultured stem cells. We further confirmed undifferentiated state (**Extended Data Fig. 5D**), isogenic derivation from the parental cell line by fingerprinting (**Extended Data Fig. 5E**), absence of chromosomal aberrations by molecular karyotyping (**Extended Data Fig. 5F**), and absence of off-target effects at the most likely sites (**Extended Data Fig. 5G**). In summary, we generated iPSC with a triple KI of AD-causing APP mutations that can be used to study AD pathogenesis in an isogenic setting.

### AD-modeling 3BTMs replicate central AD phenotypes

To analyze AD-related pathologies in our 3D model, we differentiated WT and KI iPSC into the different cell types, generated 3BTMs, and analyzed phenotype formation over time. As expected from the introduced APP mutations, we observed about 4-fold increased total Aβ levels (**Figure 5A**) as well as a 50-fold increased Aβ42:40 ratio (**Figure 5B**) in KI compared to WT cultures. We next analyzed if this increased Aβ production would lead to Aβ deposition in 3BTMs over time. We reproducibly found a progressive accumulation of Aβ in puncta and partly neurite-like structures specifically in KI cultures (**Figures 5C, D**). This could be prevented by application of a BACE inhibitor (**Figure 5E, Extended Data Fig. 6A**), confirming phenotype specificity, and demonstrating suitability of the system to test small molecule drug treatments. Interestingly, presence of iAS and iMG increased Aβ deposition in 3BTMs (**Extended Data Fig. 6B**), despite unaltered total Aβ levels and Aβ42:40 ratio (**Extended Data Fig. 6C**). As these Aβ accumulations may represent early stages of plaque formation, we analyzed if we could also detect insoluble Aβ, indicating aggregation of the peptide. We performed sequential protein extractions to analyze the insoluble fraction and indeed found insoluble Aβ42 already in 1-month-old KI cultures, and increased amounts at 6 months of age (**Figure 5F**), suggesting progressive aggregation. We then analyzed phospho-Tau (pTau) levels as a downstream pathology marker of amyloid accumulation, to check if aspects of the Aβ-Tau axis could be studied in our system. Intriguingly, we found increased levels of pTau T181, an early tauopathy marker^58^, in KI cultures at 3 months of age (**Figure 5G, Extended Data Fig. 6D**). This effect was strongest in the presence of both glial cell types, weaker if iAS were left out, and not visible without iMG (**Figure 5H, Extended Data Fig. 6D**), suggesting that iMG and their crosstalk with iAS may play an important role in the Aβ-tau axis. Moreover, pTau detected with the PHF-1 antibody was increased, which binds to misfolded and phosphorylated tau representing advanced stages of tau pathology (**Figure 5I, Extended Data Fig. 6E**).

**Figure 5.**
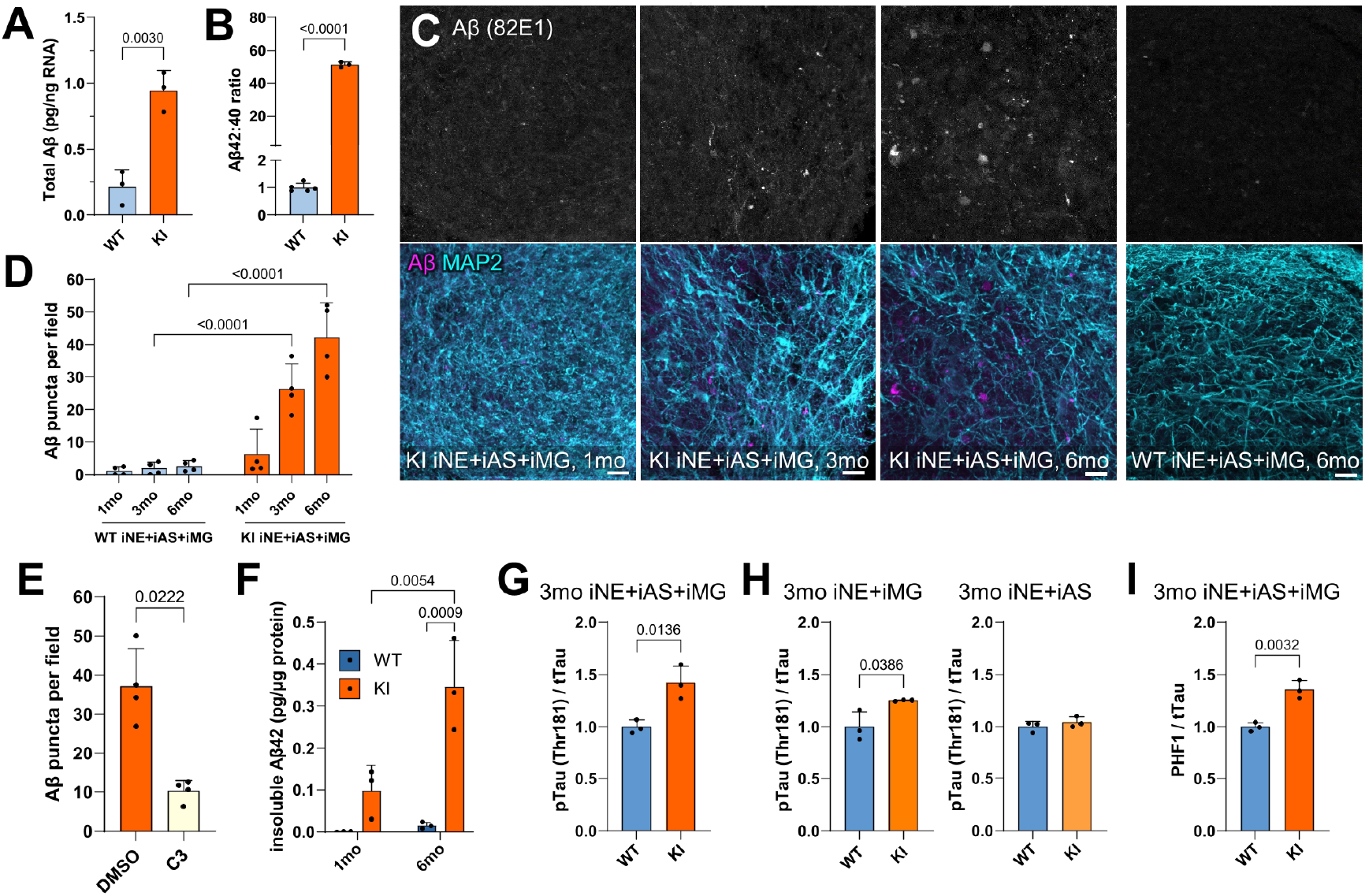
AD-modeling 3BTMs recapitulate central AD phenotypes. **(A)** MSD Immunoassay (IA) of 3BTM supernatants at 1 month to measure secretion of Aβ38/40/42 showing total Aβ levels (sum of all 3 species) (n = 3 experiments with 4-5 3BTM pooled, each dot represents one experiment, unpaired two-sided t-test). **(B)** IA as in (A) showing Aβ42:40 ratio (n = 3 experiments with 4-5 3BTMs pooled, each dot represents one experiment, unpaired two-sided t-test). **(C)** IF stainings of knock-in (KI) 3BTMs over time and WT 3BTMs at 6 months for Aβ (82E1 antibody), scale bars 20 µm. **(D)** Quantification of Aβ IF stainings as in (C) (n = 4 experiments with two 3BTMs each, each dot represents one experiment, two-way ANOVA with Šidák’s multiple comparisons test). **(E)** Quantification of Aβ IF stainings after 6-months treatment with DMSO (as control) or 5 µM BACE-inhibitor C3 (n = 4 experiments with two 3BTMs each, each dot represents one experiment, unpaired two-sided t-test), IF staining see Extended Data Fig. 6A. **(F)** IA of insoluble fractions extracted from WT and KI 3BTMs showing insoluble Aβ42 levels (n = 3 experiments with 20-25 3BTMs pooled, each dot represents one experiment, two-way ANOVA followed by Tukey’s multiple comparisons test). **(G)** Quantification of western blot (WB, see Extended Data Fig. 6D) of WT and KI iNE+iAS+iMG 3BTMs at 3 months of age detecting phospho-Tau (pTau, AT270 antibody – Thr 181 epitope) levels, normalized to total tau levels (K9JA antibody) (n = 3 experiments with 3-4 3BTMs pooled, each dot represents one experiment, unpaired two-sided t-test). **(H)** WB analysis as in (G) of 3BTMs containing iNE+iMG (left) or iNE+iAS (right) (n = 3 experiments with 3-4 3BTMs pooled, each dot represents one experiment, unpaired two-sided t-tests). **(I)** WB analysis as in (G) detecting pTau (PHF-1 antibody, Ser396/404 epitope) (for blot see Extended Data Fig. 6E) (n = 3 experiments with 3-4 3BTMs pooled, each dot represents one experiment, unpaired two-sided t-test). Data are presented as mean + SD.

### iMG in AD-modeling 3BTMs recapitulate disease-related change

Given the presence of AD-like pathology in our model, we next studied neuroinflammatory phenotypes, which is facilitated by the unique long-term presence of mature iMG and iAS in our cultures. Using qPCR, we found increased levels of *NFκB*, a central mediator of proinflammatory responses, and *CD68*, a MG activation marker (**Figure 6A**), as well as increased levels of *C3* and *C1QC* (**Figure 6B**), which have recently been implicated in inflammatory cellular crosstalk between iAS and iMG in AD^59^. To gain a comprehensive understanding of disease-associated iMG alterations in our AD model, we performed scRNAseq on KI cultures at 3 months of age, in the same experiment as the WT cultures at 1 and 3 months described above (**Extended Data Fig. 3A, B**). After subsetting our dataset for iMG from the 3-month-old WT and KI samples, we obtained 10,828 high-quality cells (**Extended Data Fig. 7A**) distributed into 6 clusters (**Figure 6C**). Again, biological replicates showed high similarity (**Extended Data Fig. 7B**), confirming reproducibility of the cultures. Strikingly, we found a major shift in iMG phenotypes comparing WT and KI, with KI iMG forming two genotype-specific clusters (*KI cluster 1* and *KI cluster 2*) distinct from the *WT resting* and the shared *chemokine, glycolytic*, and *proliferative* clusters (**Figure 6D, E**, top marker genes for each cluster are reported in **Extended Data Fig. 7C)**. We then further characterized the phenotype of the KI-specific clusters compared to the *WT resting* one. DE analysis between *KI cluster 2* and *WT resting* identified 129 significantly upregulated genes, including *TREM2, CD14*, and *PSAP*, which were previously described as upregulated in MG from human AD patients^57,58^, and 103 downregulated genes (**Figure 6F, G**). GO analysis of upregulated genes showed enrichment in pathways related to immune response and inflammation as well as chemotaxis, whereas downregulated genes were enriched in antigen presentation and processing, suggesting a shift towards a pro-inflammatory and dysfunctional phenotype in KI iMG (**Figure 6H**). Comparing *KI cluster 1* and *WT resting* yielded similar genes and terms (**Extended Data Figures 7D-F**), though with lower effect sizes, suggesting it may possess an intermediate phenotype between *WT resting* and *KI cluster 2*. We further found a significant upregulation of different GWAS-derived AD risk genes^60^, including *BLNK* and *SORL1*, specifically in *KI cluster 2* (**Figure 6I**), opening up the possibility to study the effect of these genes on MG function in the context of AD. We next compared the iMG expression signature derived from our AD model with a published MG signature from post-mortem human AD brains^61^. Intriguingly, we observed a significant enrichment of the patient-derived AD signature in iMG in *KI cluster 2* (**Figure 6K**), suggesting a replication of neuroinflammatory processes found in post-mortem AD brains in our AD model. Finally, we analyzed upregulated genes in 3BTM iMG and the published MG signature and found a large and strongly significant overlap (**Figure 6L**). Together, these findings demonstrate the suitability of our model to replicate key aspects of microglial activation seen in AD, providing a valuable tool for studying human-specific responses and early AD-related changes in iMG.

**Figure 6.**
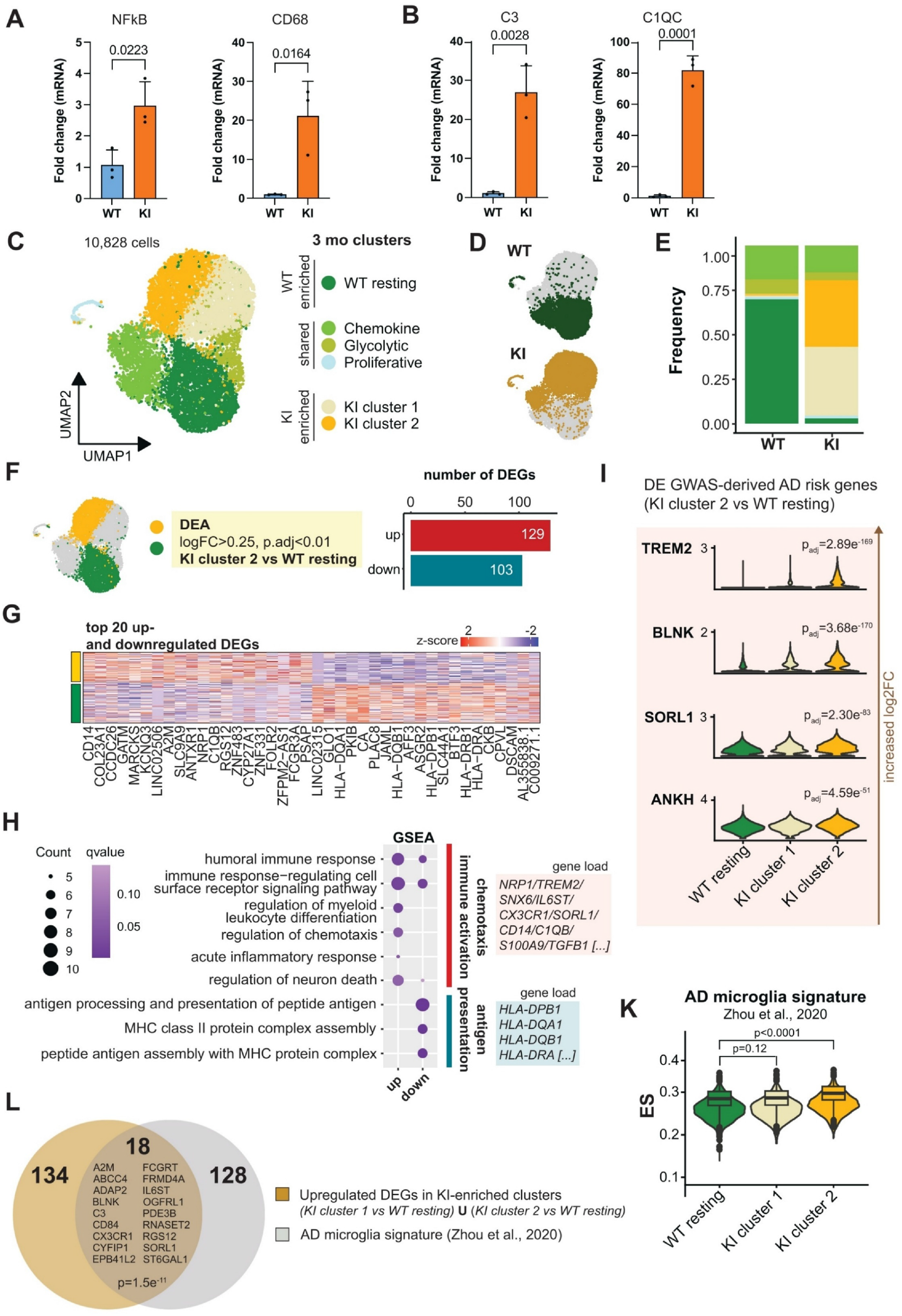
iMG in AD-modeling 3BTMs recapitulate disease-related changes. **(A)** qPCR analysis of WT and KI 3BTMs at 3 months for inflammatory markers *NFκB* and *CD68* (n = 3 experiments with 8-10 3BTMs pooled, each dot represents one experiment, unpaired two-sided t-tests). **(B)** qPCR analysis as in (A) for *C3* and *C1QC* (n = 3 experiments with 8-10 3BTMs pooled, each dot represents one experiment, unpaired two-sided t-tests). **(C)** UMAP representation (res=0.4, dims = 1:10) of the 3-month subset (subset 2, Fig. S3B) of 10,828 cells and cluster annotation of WT and KI iMG. **(D)** UMAP representation split by genotype. **(E)** Cell state frequencies reported for KI and WT genotypes separately. **(F)** Number of differentially expressed (DE) genes (p.adj<0.01, abs(log_2_FC)>0.25) obtained by comparing *KI cluster 2* and *WT resting* clusters. **(G)** Heatmap showing normalized scaled expression of the top 20 up- and downregulated DEGs. **(H)** Over-representation analysis (ORA, gene ontology, biological process) of DEGs showing enrichment of upregulated genes in chemotaxis and immune activation and of downregulated genes in antigen presentation (*q*-value<0.2). A subset of up- and downregulated genes contributing to the load of ORA functional terms is shown. **(I)** Expression level of GWAS-defined AD risk genes^67^ which are upregulated in *KI cluster 2* compared to *WT resting*, reported across *WT resting, KI cluster 1* and *KI cluster 2* cells. **(K)** Violin plot reporting the AD MG signature (by Zhou et al.^61^) enrichment score (ES) across *WT resting, KI cluster 1* and *KI cluster 2* cells (Wilcoxon signed-rank test). **(L)** Venn diagram showing overlap of KI iMG signature from this study with human AD MG signature reported by Zhou et al. (*p*-value calculated by Fisher’s exact test). Data in bar graphs ((A) and (B)) are presented as mean + SD, box plots show median, 25^th^ and 75^th^ percentiles (box) ± 1.5x IQR (whiskers).

### 3BTMs enable studying AD-relevant perturbations and drug responses in human brain cells

Given that 3BTMs replicated certain morphological and functional features of human brain tissue and allowed modulation of AD phenotypes, as we showed by BACE inhibition, we asked if the model could also be applied to investigate responses of brain cells to an AD-related perturbation and subsequent drug treatment. Anti-Aβ immunotherapy has recently been approved as the first disease-modifying treatment for AD, however, its efficiency and effects on brain function and cellular responses are still under investigation^62^. Aducanumab, one of the approved antibodies, has been shown to mostly bind fibrillar Aβ^63^. We therefore reasoned that induction of AD-like pathology with synthetic Aβ42 (sAβ42), previously shown to elicit rapid formation of fibrillar, dense-core plaques in a 2D system^5^, may be better suited for such a translational assay in 3BTMs than our genetic KI model, which shows a milder phenotype representing earlier AD stages. Dense-core plaques are a key feature of late-stage AD and have a core of highly compacted Aβ fibrils that can be stained with compounds binding to their characteristic β-sheet structure^64^, such as pFTAA^65^, and a surrounding halo of smaller, more soluble fibrils and oligomers^66^. We analyzed 3BTMs 1 month after sAβ42 treatment (at 1.5mo of age) and indeed found not only even distribution of the sAβ42 in the cultures (**Figure 7A**), but its aggregation into plaque-like structures with a core positive for pFTAA surrounded by fibrillar Aβ (**Figure 7B**), and iMG positive for Aβ (**Figure 7C**), suggesting clearance. We then treated 3BTMs after addition of exogenous sAβ42 with Aducanumab and analyzed Aβ accumulation and phagocytosis by iMG, in comparison to cultures treated with a non-targeting control antibody. After only 2 weeks of treatment, we found strong co-localization of Aducanumab but not control antibody with Aβ (**Figure 7D**), indicating successful penetration into the cultures and target engagement. Aβ load was mildly reduced over time in control-antibody treated cultures, suggesting that the peptide is cleared by iMG. Confirming efficacy of the antibody therapy, Aducanumab treatment reduced Aβ load more effectively after 2-weeks and even stronger after 1 month of treatment (**Figure 7E**). We also found colocalization of Aducanumab and Aβ inside iMG (**Figure 7F, G**), and more phagocytosed Aβ in iMG in Aducanumab-compared to control-antibody-treated cultures (**Figure 7H**), confirming the importance of iMG in the clearance process. CD68 was increased in iMG from Aducanumab-treated cultures, suggesting improved phagocytosis and activation of the cells in reaction to the treatment (**Figure 7I**). Taken together, we induced dense-core Aβ plaque-like pathology in 3BTMs by adding exogenous Aβ and showed the suitability of the model to profile anti-amyloid antibodies and microglial reaction to antibody therapy.

**Figure 7.**
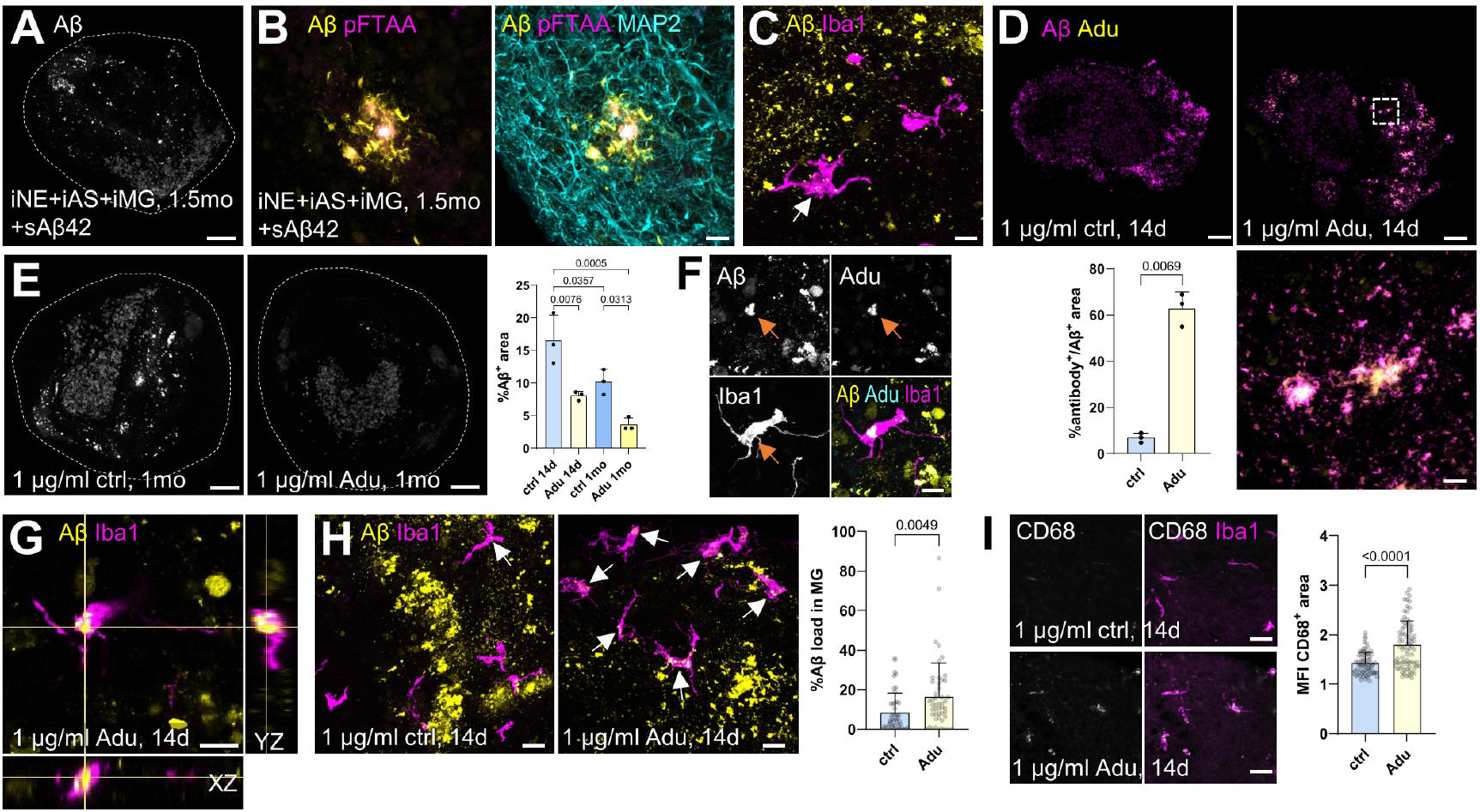
Addition of exogenous Aβ42 elicits plaque-like pathology and enables anti-amyloid antibody testing. (**A)** Aβ IF staining of iNE+iAS+iMG 3BTMs at 1.5 months of age after addition of 2x 500 nM synthetic Aβ42 on days 8 and 10, scale bar 100 µm. **(B)** Close-up of IF staining as in (A) showing plaque-like structure with aggregated core (stained by pFTAA) and fibrillar halo (positive for Aβ), scale bar 10 µm. **(C)** IF staining as in (A) showing colocalization of Aβ and Iba1 (arrow), scale bar 10 µm. **(D)** IF staining of iNE+iAS+iMG 3BTMs at 1 month after addition of 2x 500 nM synthetic Aβ42 on days 8 and 10, and treatment with 1 µg/ml control (ctrl, top left) or Aducanumab (Adu, top right) antibody on days 17, 21, 24, 28 (scale bars 100 µm). Close-up on the bottom right showing boxed region, scale bar 10 µm. Quantification in bottom left showing percentage of Aβ-positive (Aβ+) area which is also antibody-positive (n=3 experiments with two 3BTMs each, each dot represents one experiment, paired two-sided *t*-test). **(E)** IF staining as in (D) showing Aβ accumulation in cultures treated for 1 month with control antibody (left) or Aducanumab (middle), scale bars 100 µm. Quantification of Aβ-positive area after 14 days and 1 month on the right (n=3 experiments with two 3BTMs each, each dot represents one experiment, one-way ANOVA with Tukey’s multiple comparisons test). **(F)** IF staining showing co-localization of Aβ and Aducanumab inside Iba1-positive iMG, arrows showing same position in each image, scale bar 10 µm. **(G)** Orthogonal views of IF stainings shown in (F). **(H)** IF staining in 3BTMs treated as in (D), arrows pointing to colocalized signal. Scale bars 20 µm. Quantification on the right showing Aβ load (Aβ-positive area inside iMG) (n = 3 experiments with 11-22 cells per replicate and condition, each dot represents one iMG, Mann-Whitney test). **(I)** IF staining in 3BTMs treated as in (D), scale bars 20 µm. Quantification on the right showing mean fluorescent intensity (MFI) of CD68-positive area inside iMG (n = 3 experiments with 19-27 cells per replicate and condition, each dot represents one iMG, Mann-Whitney test).

## Discussion

Human 3D *in vitro* models offer great potential to study human cell biology and molecular mechanisms in health and disease. However, current brain tissue models still lack critical features that could enhance their application in basic and translational research, such as high reproducibility, long-term survival without proliferation and subsequent necrotic core formation, and efficient, long-term incorporation of homeostatic iMG. Here, we developed a new human *in vitro* model system using self-aggregation of iNE, iAS, and iMG. By first differentiating the cells in a fully defined and controlled 2D format, we avoid the variability arising from differential access to growth and patterning factors inherent to differentiation in 3D format, leading to exceptional reproducibility of the resulting cultures. Formation of 3D tissues was initiated at time points optimized for survival and postmitotic integration of all cell types, leading to reproducible culture generation across different iPSC lines approaching 100% success rate. Characterizing the 3D cultures, we show that iNE and iAS survive long-term and form dense, mature, and functional networks. iMG migrate into the cultures autonomously and develop ramifications that they use to surveil the environment, allowing quick reaction to tissue damage.

The generation of 3D *in vitro* models with lasting incorporation of “resting”, functional, and mature iMG has been a longstanding challenge. Although previous studies found limited incorporation of iMG into neural organoids^20,21,68^, these cells commonly did not show membrane expression of homeostatic marker P2RY12, were not ramified, and/or declined in number over the course of a few weeks^20,22–24^. We show that 3BTMs support prolonged maintenance of iMG for over 6 months in culture, expression of functional P2RY12 protein in the membrane, and homeostatic cell ramification, previously observed only after transplantation of iMG into mouse brains^20,48,49^. The improved survival and homeostasis of iMG in our model may be due to the absence of a necrotic core and therefore a more supportive and physiological environment compared to previous models, such as neural organoids. As organoids mimic brain development, they are characterized by precursor cell proliferation, leading to extensive growth on a millimeter scale. This can cause a hypoxic environment towards the center, and subsequent necrotic cell death, which in turn may also affect “healthy” tissue on the edge of the cultures. As MG are known to be highly sensitive to changes in their environment, this likely prevents adoption of a “resting” state. The necrosis may also contribute to iMG death or senescence due to hyperactivation, possibly explaining the rapid decline in iMG numbers in neural organoids. Another advantage of 3BTMs is the sustained presence of mature iAS in constant numbers, as confirmed by stainings and proteomics. Astrocytes have been shown to promote MG ramification and maturation^69^, and they may support the improved iMG phenotypes in 3BTMs compared to neural organoids, where immature iAS only develop after 2-3 months in culture in limited numbers^25,26^. Finally, our proteomics data indicate that 3BTMs offer a more mature tissue environment at an age relevant for iMG incorporation in comparison to organoids, possibly contributing to the more mature iMG phenotypes.

Another distinctive feature of 3BTMs is the absence of an artificial ECM during aggregation and the endogenous development of a mature, brain-like ECM. Many current *in vitro* models use Matrigel, a mouse-tumor-derived matrix, to embed cells and promote long-term stability and tissue architecture^70^. However, as a basement-membrane matrix, Matrigel is mainly composed of Laminins and Collagens^71^, whereas brain ECM mainly contains Hyaluronan, chondroitin-sulphate proteoglycans, and linker proteins^72^. As the brain ECM has been shown to influence cell functionality^11^, these differences likely impair physiological cell maturation and function. In addition, Matrigel contains various growth factors differing between lots, making it an unpredictable variable in 3D cultures. We therefore avoided the use of Matrigel or other artificial matrices in 3BTMs and instead found that the included cell types spontaneously form a mature, brain-like ECM.

Another advantage of our model is its modularity, allowing specific combinations of different cell- and genotypes for neurons, astrocytes, and microglia, which is not possible in model systems developing from the same stem or precursor cells, such as neural organoids and spheroids, where all cell types inherently share the same genetic background. In contrast, by differentiating and combining all cell types separately, our model facilitates the analysis of cell type interactions by adding or omitting cell types from the cultures. Furthermore, 3BTMs may be applied to investigate cell-type specific disease contributions of risk alleles, such as ApoE4, by analyzing cultures in which only one cell type carries the mutation. In addition, cultures representing different brain regions can be generated, e.g., hippocampus or midbrain, by simply combining iNE and glial cells pushed into the respective region-specific lineage during differentiation.

Thereby, the model may also be used to study other diseases, such as Parkinson’s disease, which strongly affects the midbrain.

Using the mature tissue environment provided by our system, we aimed to induce AD pathology by knocking in a combination of APP mutations. As shown in the corresponding mouse model^55^, the mutations have synergistic effects on Aβ aggregation, and they also induced accumulation and aggregation of Aβ in our *in vitro* system. Intriguingly, cells displayed increased tau phosphorylation using AT270 (pTau 181) and PHF-1 (pTau 396/404) antibodies, showcasing the ability to study the Aβ-tau axis, one of the most important open questions in AD research. A first set of modular experiments with glial cells in our system suggested that iMG and their crosstalk with iAS may be at the interface between the two pathologies. Due to their modularity and reproducibility, 3BTMs allow further investigating such crosstalk and elucidating the underlying mechanisms and roles of each cell type in the pathological cascade. In addition, we found evidence of neuroinflammation and a related shift in iMG states by transcriptomics, with an upregulation of AD-associated genes such as *TREM2, CD14, PSAP*, and *C1QB*. TREM2 is involved in chemotaxis towards and subsequent containment and removal of Aβ plaques ^73–75^. CD14 has previously been described as a receptor for Aβ which induces microglia activation^76,77^ and modulates AD pathogenesis^78^. PSAP is a lysosomal protein whose levels correlate with AD pathology in patients^79^. C1QB is involved in synaptic pruning^80^, a developmental process thought to be aberrantly re-activated in early stages of AD^81^. In addition, all four have been previously shown to be increased in MG from post-mortem human AD brains^61,82^. Overall, these data showcase unique opportunities to study early AD-relevant mechanisms in 3BTMs, which can hardly be studied in human brain due to inaccessibility of living brain tissue, and the fact that the processes modeled here take place years to decades before diagnosis.

In addition, due to high reproducibility and simplicity of culture generation, the model can be efficiently scaled and applied for drug testing, thus providing opportunities to overcome the scarcity of AD models replicating central disease features and being amenable for drug screening, which is a major hurdle for drug development^83^. To explore the suitability of 3BTMs for testing AD-relevant drugs, we confirmed the efficiency of β-secretase inhibitor IV (C3) in preventing Aβ accumulation in the NL-G-F knock-in model, as well as anti-amyloid antibody Aducanumab in removing aggregated Aβ after treatment with synthetic Aβ42.

The 3BTM model still has several limitations: although it enables investigation of several AD-relevant phenotypes, certain pathomechanisms are not yet fully recapitulated: formation of dense-core plaques typical for late AD stages was achieved upon treatment with sAβ42, whereas aggregation in the genetic KI model could only be detected biochemically, indicating presence of a milder phenotype more similar to early AD. This difference may be due to slower generation and build-up of pathologic Aβ in the KI 3BTMs, and its removal by degradation and media changes, which may be optimized in future work by maintaining higher Aβ levels in 3BTMs. Furthermore, although we observed a preferential localization of iMG in areas with Aβ pathology, we only saw very limited clustering of microglia around plaque-like structures in sAβ42-treated cultures. This may be explained by the lack of neuritic plaques and coinciding neurofibrillary tangle pathology, which have been shown to induce dense MG clustering in AD patient brains84. Formation of late-stage Tauopathy is limited in all iPSC-based AD models available to date, likely due to absence of adult Tau isoform expression4. In addition, Aβ clearance through the blood-brain barrier or other vascular contributions including amyloid-related imaging abnormalities (ARIA) and cerebral amyloid angiopathy (CAA), also in relation to anti-amyloid immunotherapy, cannot be investigated yet due to the absence of blood vessels. Even though the lack of these features precludes their translational investigation, effects of Aβ-targeting small molecules, antiamyloid antibodies or other therapeutics on different brain cell types can be investigated, which provides highly relevant information to improve our understanding of disease processes and develop effective therapeutic strategies.

Altogether, we developed a fully stem-cell-based human cortical tissue model that enables *in vitro* analysis of neurons and glia, including homeostatic microglia, investigating their interactions under physiological and pathological conditions, as well as applications for drug development.

## Acknowledgments

We thank Johannes Trambauer for advice on experiments and data analysis, Dennis Crusius, Caterina Mongiello, Annika Wagener, Ecem Balcioglu, Georg Kislinger, Anna Berghofer, Markus Utzt, Cornelia Niemann, and Yuxiao Zhang for technical help, Otto Windl for help with patient samples, Lorenzo Bonaguro and Simon Schäfer for help with and discussion about the scRNA sequencing analysis, and Kathryn Monroe and Joseph Lewcock for providing resources. This study was supported by Deutsche Forschungsgemeinschaft (DFG, German Research Foundation) under Germany’s Excellence Strategy within the framework of the Munich Cluster for Systems Neurology (EXC 2145 SyNergy, ID 390857198, to MS, SC, TM, NP, JH, CH, SFL, and DP), the Bonn ImmunoSensation2 cluster (EXC2151, ID 390873048, to MDB, JLS and CC), and TRR 274/1,2 project Z01 – ID 408885537 (to MS and TM). Further support came from the donors of the ADR AD2019604S (to DP), a program of the BrightFocus Foundation. This work was also funded through the Centers of Excellence in Neurodegeneration (grant CoEN6005, to SFL) and the Bundesministerium für Bildung und Forschung (BMBF, FKZ: 16LW0473, to DP; and FKZ: 01ED2402A under the aegis of JPND, to SFL and DP). This work was supported by the Ministry of Culture and Science of North Rhine-Westphalia. NGS analyses were carried out at the Joint Scientific Facility WGGC-Bonn PRECISE. Graphics from figures 1A, 1G, and 3A were created in BioRender (1A, 1G: https://BioRender.com/h8a1o1y, 3A: https://BioRender.com/i2438ox)

## Author contributions

J.K., C.C.G., and D.P. designed the study and wrote the original manuscript draft; J.K., C.C.G., S.A.M., S.F., J.J.S., L.P., B.N., A.D., C.C. and M.S. performed experiments and analyzed data, J.V.M. and C.C. analyzed the scRNA sequencing data, A.D., S.R., V.P., N.S., J.G.G., S.C., N.P., E.B., T.M., J.H., E.D.D., M.D.B., C.H., and S.F. provided essential resources for the study, N.P., E.B., J.H., C.H., S.F.L., J.L.S. and C.C. provided supervision, D.P., M.D.B., J.L.S. provided funding for the study, all authors reviewed and edited the original manuscript draft.

## Supplementary data

**Link: https://www.isd-research.de/705f5340401ade48**

**Supplementary Video 1. 3BTMs show spontaneous and synchronized activity in calcium imaging**. Whole-mount calcium imaging of 3BTM at 3 months of age using Fluo-4, single z-plane, time in min:sec, scale bar 20 µm. Related to Extended Data Fig. 4.

**Supplementary Video 2. iMG have close contact to and engulf synaptic material in 3BTMs**. 3D reconstruction of an iMG cell (Iba1, red) and all Synapsin-1 puncta (white) which colocalize with the iMG, in 3BTM at 1 month, showing close contact and engulfment of synaptic material by iMG. Related to Figure 4.

**Supplementary Video 3. iMG actively surveil their environment in 3BTMs**. 2-photon live imaging of membrane-eYFP-labeled iMG under baseline condition in iNE+iAS+iMG 3BTM at 3 months, maximum intensity projection of a 24 µm z-stack, scale bar 20 µm. Related to Figure 4.

**Supplementary Video 4. Microglia react to tissue damage in 3BTMs**. 2-photon live imaging of membrane-eYFP-labeled iMG after focal laser lesion in iNE+iAS+iMG 3BTM at 3 months, maximum intensity projection of a 22 µm z-stack, scale bar 20 µm. Related to Figure 4.

## Materials and Methods

### Human subjects

We analyzed brain tissue samples from two fetal donors (GW 15, male, unknown cause of death, post-mortem interval ∼3 days; and GW 17, male, cause of death: amniotic infection syndrome, post-mortem interval ∼5 days) and cortical tissue from the frontal pole of one juvenile donor (6 years, female, white/caucasian, post-mortem interval 11 hrs). Experiments were approved by the ethics committee of the University Hospital, LMU Munich, and informed consent for collection of tissue samples and use for research purposes was given by the legal guardians in accordance with all relevant guidelines and regulations.

### Maintenance of iPSC lines

iPSC experiments were performed in accordance with all relevant guidelines and regulations. Wildtype iPSC-lines used in this study: 1) 7889SA2, a single cell clone derived from the male 7889SA line described by Paquet et al.^56^ (RRID: CVCL_AD15, use was approved by Rockefeller University Institutional Review Board after informed consent was obtained from subjects by Coriell Institute), 2) commercially available, female iPSC line A18944 (Thermo Fisher, Cat#A18945), and 3) commercially available, male iPSC line GM05399 (Coriell)., RRID: CVCL_7425). iPSC were maintained as described previously^29^ on vitronectin (VTN, Thermo Fisher, Cat#A14700)-coated plates in Essential 8 Flex medium (E8F, Thermo Fisher, Cat#A2858501) in a humidified incubator at 37 °C, 80-90 % humidity, 5% CO_2_, and ambient O_2_ levels. Cells were kept as colonies and split every 3-4 days using PBS (Thermo Fisher, Cat#14190094) with 0.5 mM UltraPure EDTA (Thermo Fisher, Cat#15575020). iPSC lines are available from the authors after filling out an MTA.

### Cortical neuron differentiation from iPSC

Cortical neuron differentiation was performed as described by us recently^29^. Briefly, single iPSCs were split confluently into wells of a 12-well plate coated with Geltrex (Thermo Fisher, Cat#A1413302) and neural lineage was induced using dual-SMAD inhibition by adding neural differentiation medium (Neurobasal (Cat#21103-049) and DMEM/F12 (Cat#11320-074) in a 1:1 ratio with 1x B27 supplement (Cat#17504001), 1x NEAA (Cat#11140-050), 0.5x N2 supplement (Cat#17502048), 1x GlutaMAX (Cat#35050), 1x Penicillin/Streptomycin (Cat#15140-122), 50 µM β-mercaptoethanol (Cat#21985-023, all Thermo Fisher), and 2.5µg/ml Insulin (Sigma-Aldrich, Cat#I0516)) supplemented with 10 µM SB431542 (Selleckchem, Cat#S1067) and 0.25 µM LDN-193189 (Selleckchem, Cat#S2618) for 10 days, WNT inhibitor XAV939 (5 µM, Selleckchem, Cat# S1180) for the first 2 days, and ROCK-inhibitor (10 µM Y27632, Selleckchem, Cat#S1049) for the first day. Neural rosettes were expanded on plates coated with poly-L-ornithin (Sigma-Aldrich, Cat#P4957) and Laminin (Thermo Fisher, Cat#23017015) by addition of 20 ng/µl bFGF (STEMCELL Technologies, Cat#78003.2) upon formation, and isolated manually using Neural Rosette Selection Reagent (STEMCELL Technologies, Cat#05832) around day 20-23. Rosettes were further expanded in neural differentiation medium for 7 days, then split using Accutase (Sigma-Aldrich, Cat#A6964) and grown for another 7 days, before being frozen in neural induction medium supplemented with 10% DMSO (Sigma-Aldrich, Cat#D2650) and 20 ng/µl bFGF. After thawing, rosettes were further differentiated for 5-7 days, and then used for 3BTMs on day 38-43 of differentiation.

### Astrocyte differentiation from iPSC

Differentiation of iPSC to astrocytes was performed similar to the protocol described by Perriot et al.^30,85^, with modifications. Briefly, iPSC were differentiated as in the cortical neuron differentiation protocol for 13-15 days, when rosettes became visible. Media was then changed to glial precursor expansion medium (DMEM/F12 with 1x B27 supplement minus vitamin A (Thermo Fisher, Cat#12587010), 1x N2 supplement, 1x GlutaMAX, 10 ng/ml bFGF, 10 ng/ml EGF (Peprotech, Cat#100-15), and rosettes were manually isolated after 3 days as described in the neuron differentiation protocol. After growing confluent, rosettes were split to single cells using Accutase after which the protocol by Perriot et al. was followed. Glial expansion medium was applied for a total of 6-9 days, astrocyte induction medium (DMEM/F12 with 1x B27 minus vitamin A, 1x N2, 1x GlutaMAX, 10 ng/ml EGF, 10 ng/ml LIF (Peprotech, Cat#300-05)) for 14 days, and astrocyte medium (DMEM/F12 with 1x B27 minus vitamin A, 1x GlutaMAX, 20 ng/ml CNTF (Peprotech, Cat#450-13) for 28 days, after which cells were frozen in astrocyte medium supplemented with 10% DMSO. After thawing, astrocytes were grown for 5-7 more days and then used for 3BTMs on day 64-69 of differentiation.

### Microglia differentiation from iPSC

Differentiation of iPSC to microglia was performed following the protocol described by us^31^ and others^86^ recently. Briefly, 80-90% confluent iPSC colonies were split 1:150-1:200 using PBS with EDTA, and hematopoiesis was induced for 12 days using the STEMDiff Hematopoietic Kit (STEMCELL Technologies, Cat#05310) following the manufacturer’s instructions. Floating hematopoietic precursor cells were collected and frozen in Bambanker (FujiFilm Wako, Cat#302-14681). After thawing, hematopoietic precursor cells were differentiated into microglia for 12 days in DMEM/F12 supplemented with 2x Insulin-Transferrin-Selen (ITS-G, Thermo Fisher, Cat#41400-045), 2x B27 supplement, 0.5x N2 supplement, 1x GlutaMAX, 1x NEAA, 5 µg/ml Insulin, 1x Penicillin/Streptomycin, and 400 µM monothioglycerol (Sigma-Aldrich, Cat#M1753), 100 ng/ml IL34 (Peprotech, Cat#200-34), 50 ng/ml TGF-β1 (Peprotech, Cat#100-21), and 25 ng/ml M-CSF (Peprotech, Cat#300-25). Microglia were used for 3BTMs on day 24 of differentiation.

### Generation of 3D cortical brain tissue models (3BTMs)

To prepare 3BTMs only containing neurons, rosettes/early neurons were split into single cells using Accutase, and plated into ultra-low-attachment, 96-well round-bottom plates (Sigma-Aldrich, Cat#CLS7007) at 250,000 cells/well in 3D culture maintenance media (Neurobasal Plus (Thermo Fisher, Cat#A3582901) with 1x B27 Plus supplement (Thermo Fisher, Cat#A3582801), 0.25x GlutaMAX, and 1x Penicillin/Streptomycin). Cells were spun down for 2 min at 200 *g* and cultured at 37 °C, 5% CO_2_, 80-90% humidity and ambient oxygen levels. 3D culture maintenance media was supplemented with 10 μM DAPT (Selleckchem, Cat#S2215) for 7 days after culture generation, and with 5 μM 5-Fluorouracil (5-FU, Sigma-Aldrich, Cat#F6627) and 5 μM Uridine (Sigma-Aldrich, Cat#U3750) from day 3 to day 15 after culture generation. 7-10 days after generation, aggregated 3D cultures were transferred to wells of a 12-well plate coated with anti-adherence rinsing solution (StemCell Technologies, Cat#07010) for long-term culture with half of the medium being replaced every 3-4 days. To prepare 3BTMs containing neurons and astrocytes, cells were split as described above, and plated at a neurons:astrocytes = 3:1 ratio with a total of 250,000 cells, meaning 187,500 neurons and 62,500 astrocytes. To prepare 3BTMs containing neurons, astrocytes, and microglia, neurons and astrocytes were mixed and aggregated as described above. On day 15 after culture generation, when treatment with 5FU and uridine ended, cultures were transferred to ultra-low attachment, 96-well V-bottom plates (Greiner Bio-One, Cat#651970) in fresh 3D culture maintenance medium supplemented with 100 ng/ml IL43, 50 ng/ml TGF-β1, and 25 ng/ml M-CSF. Microglia were collected after differentiation and added into the wells at 37,500 cells per 3D culture resulting in an initial neuron:microglia ratio of 5:1. When microglia had migrated into the cultures (7-10 days), cultures were transferred back to 12-well plates coated with anti-adherence rinsing solution for long-term culture as described above with continuous addition of IL34, TGF-β1, and M-CSF. 3BTMs were generated from lines 7889SA2 and A18944, and from the isogenic APP KI line corresponding to 7889SA2.

### Generation of neural organoids (NOs)

Neural organoids were generated according to Lancaster et al.^14,15^ with minor modifications. Briefly, iPSC were dissociated into single cells using Accutase, and 9,000 cells per well were seeded to an ultra-low attachment, round-bottom 96-well plate in human stem cell media (75% DMEM-F12, 20% knock-out serum replacement (Thermo Fisher, Cat#10828028), 3% FBS (Thermo Fisher, Cat#10270106), 1% GlutaMAX, 1% MEM-NEAA, 50 µM β-mercaptoethanol) with 4 ng/ml bFGF and 50 μM ROCK-inhibitor. After 6 days, media was replaced for neural induction media (DMEM-F12 with 1x N2 supplement, 1x GlutaMAX supplement, 1x NEAA, 1 µg/ml heparin (Sigma-Aldrich, Cat#H3149), and neuroepithelial tissues were fed every other day for 6 days. On day 12, embryoid bodies were transferred to droplets of Matrigel (Corning, Cat#356230) in differentiation media (Neurobasal and DMEM/F12 in a 1:1 ratio with 1x B27 supplement minus vitamin A, 0.5x NEAA, 0.5x N2 supplement, 1x GlutaMAX, 2.5 µg/ml insulin, 1x Penicillin/Streptomycin, 50 µM β-mercaptoethanol). After 4 days of stationary growth, organoids were moved to an orbital shaker at 57 rpm, 37°C, 5% CO_2,_ and ambient oxygen levels in differentiation medium with vitamin A. Media was then replaced every 3-4 days. On Day 90, organoids were collected and washed with PBS. Once all the liquid was removed, organoids were snap-frozen on dry ice and kept at -80 °C. Neural organoids were generated from line GM05399.

### Immunofluorescence stainings

For 2D immunofluorescence staining, cells were plated on acid-etched coverslips (Marienfeld, Cat#630-2190) as described before^29^. At the day of fixation, cells were washed once with PBS and fixed with 4% PFA (Morphisto, Cat#1.030.300.250) for 10 min at RT. Cells were then washed with PBS for 3× 5 min and blocked for 1 hour at room temperature (RT) in blocking solution (PBS with 3% donkey serum (Sigma-Aldrich, Cat#D9663), 0.1% Triton X-100 (Sigma-Aldrich, Cat#T8787) and 0.02% NaN_3_ (Sigma-Aldrich, Cat#S2002) in PBS. Primary antibodies were diluted in blocking solution and incubation was performed overnight at 4 °C. Coverslips were washed with PBS for 3×5 min and then incubated with secondary antibodies (1:500 dilution), and DAPI if required (Thermo Fisher, Cat#D1306, stored as 5 mg/ml stock solution, 1:50,000 dilution), diluted in blocking solution for 2 hours at RT in the dark. Cells were washed again for 3x 5 min with PBS and mounted on glass slides (Epredia, Cat#J1800AMNZ) in Fluoromount-G (Thermo Fisher, Cat#00-4958-02). Coverslips were imaged with an inverted fluorescent microscope (Zeiss Axio Observer Z.1) and images were processed in Fiji^87^.

For 3D culture stainings, cultures were washed once with PBS, fixed with cold 4 % PFA for 30 min at RT, and washed again with PBS for 3x 30 min. Cultures were then transferred into 30 % sucrose (Sigma-Aldrich, Cat#S0389) in PBS over night at 4 °C, then transferred into OCT compound (Sakura Finetek Tissue-Tek O.C.T. Compound, Fisher Scientific, Cat#1235175), and cryosectioned into 30 µm slices. Slices were mounted onto glass slides, dried on a heat plate at 45°C, rehydrated with H_2_O for 3 min, and transferred into a humidified box. All following steps starting with the blocking were performed as described above, with 3x10 min washes in between incubations. TUNEL stainings were performed on slides as described above using the In Situ Cell Death Detection Kit, TMR red (Sigma-Aldrich, Cat#12156792910) according to manufacturer’s instructions. Hyaluronan (hyaluronic acid) was stained using biotin-coupled hyaluronic-acid-binding-protein (Hölzel Biotech, Cat#HKD-BC41) followed by avidin-FITC-based detection (Thermo Fisher, Cat#434411) following the protocol above. For hypoxia stainings, 3BTMs were incubated with 200 µM Pimonidazole HCl (Hypoxyprobe Omni Kit, Hypoxyprobe, Cat#HP3-100Kit), to label proteins in hypoxic cells^88^, for 2h at standard culture conditions (ambient O_2_ levels), or at 5% O_2_ for positive controls. Stainings were performed according to the manufacturer’s instructions, with primary antibody dilution 1:50. Samples were imaged using a Zeiss LSM 880 inverted confocal microscope using 10x (for overview images, EC Plan-Neofluar/0.3 Air), 40x (for close-ups, EC Plan-Neofluar/1.30 Oil) and 100x objectives (for synapses, alpha Plan-Apochromat/1.46 Oil) on ZEN black software (Version 10.0.4.910), and images were processed and analyzed with Fiji. Images shown are representative maximum-intensity projections of z-stacks acquired of the 30 µm slices. All quantifications were done on unedited, raw images. TUNEL-positive cells, Ki67-positive cells, NeuN/Sox9-positive cells, DAPI-positive cells, Aβ puncta, Aβ area after seeding and antibody treatment, and CD68 signal inside microglia were quantified by thresholding the relevant channel in the raw images and subsequent use of the “Analyze Particles” function or measuring intensity in the positive area (for seeding and antibody treatment experiments). Microglia morphology was assessed using the “MotiQ” plugin^88^ by cropping each microglia cell manually, thresholding with the “Li” algorithm and using the 3D analyzer with default settings. The ramification index is defined as the ratio of the cell surface area to the surface area of a sphere with the same volume as the respective cell, whereas the number of branches is quantified based on a skeleton of the cell created by the plug-in. Synapses were quantified using the SynQuant plugin^89^ with default setting. 3D surface renderings of synapses on/in microglia were done using Imaris Version 9.4 (Oxford Instruments). First, a surface was created based on the Iba1 staining to have a representation of the microglia, then only synapsin-1-positive puncta touching this surface were displayed to remove all synapses not in contact with/not inside the microglia. For IF stainings, if not indicated otherwise, 1-2 cultures per experiment with 2-3 slices per 3BTM were analyzed as technical replicates and pooled for analysis.

### Mass spectrometry

#### Sample preparation

3BTMs, neural organoids, or human fetal and juvenile brain samples were washed once with PBS and dissociated in RIPA buffer (50 mM HEPES pH 7.5 (Sigma-Aldrich, Cat#SRE0065), 500 mM LiCl (Sigma-Aldrich, Cat# L9650), 1 mM EDTA pH 8, 1% NP-40 (Sima-Aldrich, Cat#56741), 0.7 % sodium deoxycholate (Sigma-Aldrich, Cat#30970-25G) supplemented with protease inhibitors (Sigma-Aldrich, Cat#4693159001) and phosphatase inhibitors (Sigma-Aldrich, Cat#4906845001) using ceramic beads (Precellys Ceramic kit, VWR, Cat#432-0293) on a Precellys Evolution homogenizer at 6500 rpm for 30 sec at 4 °C. Three 3BTMs were pooled for each sample to increase total protein yield, and from fetal brain sample 1 (GW 17), two pieces of tissue were extracted. Lysates were transferred to protein low-binding tubes and spun down for 20 min at 18,000 *g* and 4 °C. Protein concentrations of supernatants were measured, and samples were frozen at -80 °C until analysis. 15 µg per sample were subjected to tryptic digestion. To this end, MgCl_2_ (10 mM) was added, and DNA was digested with 25 units Benzonase (Sigma-Aldrich, Cat#E1014) for 30 min at 37°C. Proteins were reduced using 15 mM dithiothreitol (DTT) for 30 min at 37°C, followed by cysteine alkylation with 60 mM iodoacetamide (IAA) for 30 min at 20 °C. Excess IAA was removed by adding DTT. Detergent removal and subsequent digestion with 0.2 µg LysC and 0.2 µg trypsin (Promega) was performed using the single-pot, solid-phase-enhanced sample preparation as previously described^90^. After vacuum centrifugation, peptides were dissolved in 20 µL 0.1% formic acid (Sigma-Aldrich) and peptide concentrations estimated using the Qubit protein assay (Thermo Fisher).

#### LC-MS/MS analysis

350 ng of peptides were separated on a nanoElute nanoHPLC system (Bruker, Germany) in an in-house packed C18 analytical column (15 cm × 75 µm ID, ReproSil-Pur 120 C18-AQ, 1.9 µm, Dr. Maisch GmbH). Peptides were separated with a binary gradient of water and acetonitrile (B) containing 0.1% formic acid at flow rate of 300 nL/min (0 min, 2% B; 2 min, 5% B; 94 min, 24% B; 112 min, 35% B; 120 min, 60% B) and a column temperature of 50°C.

The nanoHPLC was online coupled to a TimsTOF pro mass spectrometer (Bruker, Germany) with a CaptiveSpray ion source (Bruker, Germany). For relative protein quantification, a Data Independent Acquisition (DIA) – Parallel Accumulation Serial Fragmentation (PASEF) method was used^91^. Each scan cycle included one full MS scan followed by 34 windows of 26 m/z width (1 m/z overlap) covering an m/z range of 350-1200 using 2 windows per PASEF ramp time of 100 ms. This resulted in a cycle time of 1.9 s. Neural organoid and juvenile brain samples were run in three technical replicates.

#### Data Analysis

The MS raw data was analyzed with DIA-NN software (Version 1.8)^92^ using a library-free search. Trypsin was defined as protease and 2 missed cleavages were allowed. The data was searched against a canonical one protein per gene human protein database from UniProt (download: 2023-03-01, 20603 entries) including a fasta database with common contaminants (246 entries) from MaxQuant^92^. Oxidation of methionines and acetylation of protein N-termini were defined as variable modifications, whereas carbamidomethylation of cysteines was defined as fixed modification. The precursor and fragment ion m/z ranges were limited from 350 to 1200 and 200 to 1700, respectively. Precursor charge states were set to a range of 2-4. Peptide and peptide fragment tolerances were optimized by DIA-NN. The match between runs option was enabled. An FDR threshold of 1% was applied for peptide and protein identifications. The generated spectral library includes 13552 protein isoforms, 15198 protein groups and 222055 precursors in 180693 elution groups. Afterwards, all samples were searched against this spectral library for protein label-free quantification (LFQ). Identified proteins of the contaminants database and keratins were deleted from the results tables. One sample (sample 24) was excluded as it did not meet quality control standards. For statistical analysis, only proteins were considered which were quantified in all replicates of at least one experimental group. After log_2_ transformation of LFQ intensities, missing log_2_ LFQ intensities were imputed from a normal distribution using a width of 0.3 standard deviations of the data with a downshift of 1.8 using the software Perseus (Version 1.6.14.00)^93^. PCA analysis was performed on imputed log_2_ LFQ values without scaling or centering. GO analysis (GO_BP_direct) was performed with DAVID^94,95^ using the top/bottom 400 proteins from PCA loadings sorted by the relevant PC, only peptides with one unique UniProt AC were used. Heatmaps were generated from imputed log_2_-fold-changes of LFQ values of the different samples in comparison to the average of log_2_ LFQ values of the 3-day-old iNE+iAS samples as baseline. Hierarchical clustering was performed using Euclidean distance and average clustering. Synapse-associated proteins are all proteins within the GO terms “synapse” (GO:0045202), “transmission of nerve signals” (GO:0019226), or “neurotransmitter transport” (GO:0006836) downloaded from www.uniprot.org in December 2023 that were detected in any of the samples. Extracellular matrix proteins are a manually curated list of known human brain ECM proteins that were detected in any of the samples. Astrocyte maturation markers were curated from a list of mature AS marker genes published by Zhang et al.^37^. Corresponding proteins that were found in any of the samples were then filtered for AS-specific expression to avoid confounding effects of expression in neurons and/or microglia. For filtering, human expression data across brain cell types was downloaded from www.brainrnaseq.org, and only proteins whose gene expression in iAS was more than 10x higher than in iNE and more than 3x higher than in iMG were included in the final list.

Cell cycle markers are a manually curated list of known cell-cycle associated proteins found in the dataset.

### Transmission Electron Microscopy (TEM)

3BTMs were immersion fixed (4% PFA and 2.5% glutaraldehyde in 0.1 M sodium cacodylate buffer, pH 7.4; Science Services) for 24h. A standard rOTO en bloc staining protocol^96^ was applied including postfixation in 2% osmium tetroxide (EMS), 1.5% potassium ferricyanide (Sigma) in 0.1 M sodium cacodylate (Science Services) buffer (pH 7.4). The staining was enhanced by reaction with 1% thiocarbohydrazide (Sigma) for 45 min at 40°C. 3BTMs were washed in water and incubated in 2% aqueous osmium tetroxide, washed, and then further contrasted by overnight incubation in 1% aqueous uranyl acetate at 4°C and 2h at 50°C. Samples were dehydrated in an ascending ethanol series and infiltrated with LX112 (LADD). Cured blocks were trimmed on an EM TRIM2 (Leica) instrument to expose the center of the spheroid. Ultrathin sections were generated using a diamond knife (Diatome) on a UC7 microtome (Leica) and deposited onto formvar-coated copper grids (Science Services). Sections were imaged using a JEM-1400+ (JEOL) equipped with a XF416 (TVIPS) and the EM-Menu software (TVIPS). Images analysis was performed in Fiji.

### Single-cell RNA sequencing (scRNAseq)

To obtain a single cell suspension for scRNAseq, 4-5 cultures from 2 independent experiments (=biological replicates) at 1 or 3 months of age were dissociated using the Neural Tissue Dissociation Kit P (Miltenyi Biotec, Cat#130-092-628) in a mix of 2.375 ml buffer X, 62.5 µl enzyme P, 10 µl enzyme A, 20 µl buffer Y and 11.25 µM Actinomycin D (Sigma-Aldrich, Cat#A1410) on a gentleMACS Octo Dissociator (Miltenyi Biotec). A custom dissociation protocol was used (+ means counter-clockwise rotation, - means clockwise rotation): 4.5min +20rpm, 0.5min -100rpm, 4.5min +20rpm, 0.5min - 250rpm, 4.5min +20rpm, 0.5min -300rpm, 4.5min +20rpm, 0.5min -300rpm, 2min +20rpm. The dissociated cell suspension was continuously kept on ice or at 4 °C, and plasticware was pre-treated with 1 % bovine serum albumin solution (BSA, Thermo Fisher, Cat#15260-037) in PBS to prevent cell adhesion. Cells were filtered through a 70 µm cell strainer to remove residual clumps, spun down at 300 *g* for 5 min, and resuspended in 50 µl FACS buffer (Thermo Fisher, Cat#00-4222-57). For multiplexing of biological replicates, cell suspensions were incubated for 20 min with Hashtag-coupled Antibodies (Biolegend), washed once with FACS buffer, pelleted for 5 min at 300 *g*, and resuspended in 300 µl FACS buffer. Single cells were sorted on a FACSAria Fusion (BD Biosciences) into cooled DNA low-binding tubes (Eppendorf), DAPI (250 ng/ml) was added 5 min before sorting to exclude dead cells. Identical cell numbers of biological replicates (labelled with hashtag antibodies) were pooled into one collection tube, washed once in 0.04% BSA/PBS, and ∼16,500 cells per sample were loaded onto a 10x Chip G (10× Genomics). Single cell gene expression and cell hashing libraries were generated on the 10x Genomics Chromium platform using the Chromium Next GEM Single Cell 3’ Reagent Kits v3.1 according to the manufacturer’s protocol (CG000317 Rev C). Gene expression and hashtag libraries were sequenced on a NovaSeq 6000 S4 v1.5 flow cell (Illumina).

All analyses were performed in a containerized environment created expanding the docker image available at: https://github.com/jsschrepping/r_docker to ensure reproducibility. For the pre-processing of the Genomics GEX and TotalSeq data, 10x Genomics Cell Ranger v7.1.0^97^ was used with default parameters. The raw sequencing reads were aligned against the default genome reference “refdata-gex-GRCh38-2020-A” using the function “cellranger multi”, and the data was demultiplexed by sample origin employing the antibody tag sequences for the TotalSeq antibodies (Biolegend).

Further downstream analysis was carried out with the Seurat pipeline (v4.3.0.1)^98^. A starting Seurat object was created by incorporating all data and metadata available for both 1mo WT, 3mo WT and 3mo KI samples. 16,233 cells were included in the analysis after an initial filtering to exclude lower quality cells (> 250 genes per cell; > 15,000 UMIs per cell; mitochondrial reads % < 7%, and < 25% ribosomal reads per cell, see Extended Data Fig. 3A). PCA was performed after normalization and scaling using the standard Seurat workflow. The first 10 principal components were chosen to calculate UMAP embedding adopting a clustering resolution of 0.5. Cell type identity was attributed to each cluster based on the expression of canonical markers of microglia (TYROBP), neurons (SOX11) and astrocytes (ALDH1A1, AQP4). Only microglia cells (15.479 cells) were selected for downstream analysis, as reported in Extended Data Fig. 3B. Then, two distinct subsets were analyzed (Extended Data Fig. 3B): subset 1 containing WT cells (including 1mo and 3mo cells), and subset 2 containing cells from 3-months-old 3BTMs (including KI and WT cells). For each subset, PCA and UMAP were calculated taking into consideration the first ten principal components and a clustering resolution of 0.3 (WT subset, 8,827 cells, Figure 3B) and 0.4 (3 months subset, 10,828 cells, Figure 6C) respectively. QC was performed across obtained clusters ensuring no QC influence on the final clustering. In most of the cases, clusters annotation was assigned based on the top ten markers per cluster inferred using the Seurat “FindAllMarkers” function (only.pos = TRUE, min.pct = 0.25, logfc.threshold = 0.25), as described in the Results section. Relative cell abundances were calculated with the “*propeller*” function from speckle (v1.0.0) R package^99^.

The expression of selected genes was carried out using the FeaturePlot or VlnPlot Seurat functions, reporting the normalized counts per cell for genes of interest. Further visualizations were obtained with “*Stacked_VlnPlot*” (scCustomize v1.1.1^100^). All DEAs were run using the Seurat “FindMarkers” (min.pct = 0.25, test.use= “MAST”) function. Genes with p.adj < 0.01 and log(FC) > 0.25 (upregulated) or < -0.25 (downregulated) were defined as DEGs. The “enrichGO” function was used for gene set enrichment analyses based on the Gene Ontology database (*biological process* terms, q.value < 0.2)^101^. Heatmaps of the top 20 up- and downregulated differentially expressed genes were realized with the function DoHeatmap from Seurat. Multiple signatures were employed to evaluate over-representation of genes related to different microglia maturation stages, different in vitro models and postmortem human brain affected by AD. The gene lists used as signatures were derived from Supplementary tables of the original publication or derived after dataset re-analysis (for Popova et al., 2021 data^21^). All reported signatures were scored using the function “*AddModuleScore_UCell*” from the R package UCell (v.2.6.2)^102^. The statistical significance in the overlap between upregulated genes in KI clusters and the Zhou et al. dataset^61^ was determined by Fisher’s exact test by implementing the “testGeneOverlap”^103^ and “supertest”^104^ functions. The expression of a list containing risk genes for Alzheimer’s diseases from GWAS studies curated by Mancuso et al., 2024^60^ was evaluated in this dataset and displayed using Seurat VlnPlot function to visualize the expression of the AD risk genes also found among the differentially expressed genes when comparing the clusters WT Resting 3mo and KI cluster 2.

Monocle 3 (v1.3.1)^105–107^ was used to determine trajectories and pseudotime in Figure 3I employing the functions learn_graph (use_partition = FALSE) and order_cells (root_pr_nodes = ‘Y_24’). Integration between our 1mo and 3mo WT microglia and the Han et al. scRNA-seq dataset^43^ was performed using the Seurat RPCA approach (function “IntegrateData”, k.weight = 45) following random subsampling (n = 8,827) to match cell numbers in the two datasets. For the Popova et al. *in vitro* models dataset^21^, we first obtained separate Seurat objects and integrated them using the Seurat CCA approach (“IntegrateData” function with default parameters) after SCT normalization. The datasets contained fetal-derived human primary microglia 2D-cultured in vitro (referred as 2D in the original dataset) or MG incorporated into neural organoids (oMG), as well as human iPSCs-derived microglia either cultured in 2D (called iMG in the original dataset) or mouse xenotransplanted (xMG). In vitro model-specific marker genes from the comparison of multiple in vitro datasets were determined with the function *FindAllMarkers* (only.pos = TRUE, min.pct = 0.25) by contrasting cells from each model against the others. These genes were further selected by adjusted p-value < 0.01 and fold change > 1 and the top 400-fold-change features were used to generate each model-specific signatures.

### Electrophysiological measurements

3D cultures were embedded into 4% low-melting-point agarose (Thermo Fisher). The agarose block was put to 4°C immediately after to accelerate solidification. The block was then rapidly glued on a vibratome specimen holder and placed in 2-4°C cutting solution containing NMDG, KCl (2.5 mM), MgCl_2_ (2 mM), CaCl_2_ (2 mM), NaH_2_PO_4_ (1.2 mM), HEPES (10 mM), NaHCO_3_ (21 mM), and Glucose (5 mM), adjusted to pH 7.2 with HCl. 250 µm sections were recovered in 36°C cutting solution for 15min and 40-60 minutes at RT in aCSF containing NaCl (125 mM), KCl (2.5 mM), MgCl_2_ (2 mM), CaCl_2_ (2 mM), NaH_2_PO_4_ (1.2 mM), NaHCO_3_ (21 mM), HEPES (10 mM), and Glucose (5 mM) adjusted to pH 7.2 with NaOH. Sections were continuously superfused with carbogenated aCSF while recording at a flow rate of ∼2 ml/min at 30-32°C.

Current-clamp recordings were done in perforated patch clamp configuration. Cells were visualized with a fixed-stage microscope (BX51WI, Olympus, Hamburg, Germany) using a 60x water-immersion objective (LUMplan FL/N 60x, Olympus). Electrodes were prepared from borosilicate glass (Science Products #GB150-8P) with a vertical pipette puller (PC-10; Narishige) with tip resistances between 4 and 6 MΩ. Recordings were performed with an EPC10 patch-clamp amplifier (HEKA) controlled by PatchMaster (version 2.32; HEKA), and data acquired with a micro1410 data acquisition interface and Spike 2 (version 10) (both from CED) at 25 kHz and low-pass-filtered at 2 kHz with a four-pole Bessel filter. The calculated liquid junction potential of 14.6 mV was compensated or subtracted offline (calculated with Patcher’s Power Tools plug-in from https://www.mpibpc.mpg.de/groups/neher/index.php?page=software for IGOR Pro 6 (Wavemetrics). The pipette solution used for recordings contained K-gluconate (147 mM), KCl (10 mM), HEPES (10 mM), EGTA (0.1 mM), and MgCl_2_ (2 mM) adjusted to pH 7.2 with KOH. The patch pipette was tip filled and back filled with internal solution containing the ionophore amphotericin B (200 µg/ml, Sigma-Aldrich) and 0.02 % tetramethylrhodamine-dextran (3000MW, Thermo Fisher). Access resistance (Ra) was constantly monitored during the perforation process, and experiments were started after Ra had reached a steady state (∼5–20 min) or the action potential amplitude was stable. Intrinsic characteristics were determined using a set of depolarizing and hyperpolarizing current injections either in perforated mode with stable action potential amplitude and/or membrane potential or rapidly after the membrane ruptured spontaneously.

### RNA extraction and qPCR

RNA was extracted using the NucleoSpin RNA/Protein kit (Macherey Nagel, Cat#740933.250) according to manufacturer’s instructions for cultured cells and tissues. RNA was converted into cDNA using the High-Capacity cDNA Reverse Transcription Kit (Thermo Fisher, Cat#4368814) following the manufacturer’s instructions. qPCR was performed using PowerTrack SYBR Green Mastermix (Thermo Fisher, Cat#A46109) as suggested by the manufacturer.

### 2-Photon live cell imaging

Experiments were performed using a Zeiss LSM 7 MP multi-photon microscope equipped with a Ti:Sapphire Laser (Chameleon Vision S, Coherent) and a W Plan-Apochromat 20x/1.0 DIC D=0.17 M27 75mm objective (Zeiss). Cultures were immobilized in a 60 mm dish (VWR) in 5 µl droplets of 5% low-melting point agarose and 3D culture maintenance media was carefully added. Dishes were placed on a 37°C heat pad under the microscope and imaging was done at an excitation wavelength of 950 nm. Time lapse experiments of z-stacks were done at 1 µm axial step size, 512x512 pixel frame size, 3x zoom for 10-15 minutes with 40-50 seconds per z-stack, and analyzed in Fiji. For focal laser injuries, the laser was tuned to 750 nm, and 1 frame exposure with 30% total laser power at 40x zoom was used to damage an area of ∼9 µm diameter and ∼1 µm axial extent.

### Calcium imaging

3BTMs at 3 months of age were loaded for 1h with 2.5 µM Fluo-4 AM (Biomol, Cat#ABD-20552, 1:2000 from a 5 mM stock in DMSO) in 3D culture maintenance medium with 1:2000 Pluronic F-127 (Thermo Fisher, Cat#P3000MP) to improve uptake. After 1h, cultures were washed 2x with 3D culture maintenance medium, transferred to 8-well imaging slides (µ-Slide 8 Well, Ibidi) in 3D culture maintenance medium, and imaged on a Zeiss LSM880 inverted confocal microscope using a water immersion objective (LD LCI Plan-Apochromat 25x/0.8) with 2x zoom and maximum speed (acquisition time ∼400 ms per frame). Acquisitions were analyzed in Fiji using the multi-measure function, results were exported to Excel and analyzed there.

### CRISPR/Cas9 gene editing

The CRISPOR online tool (http://crispor.tefor.net/)^108^ was used to design sgRNAs and determine the most likely off-target loci. Gene editing was performed as described previously^109,110^ with minor modifications. Briefly, sgRNAs were cloned into plasmid MLM3636 (a gift from K. Joung, Addgene #43860), and plasmid pSpCas9(BB)-2A-Puro (PX459) V2.0 (a gift from F. Zhang, Addgene #62988) was used for Cas9 expression. Single-stranded oligodeoxynucleotides (ssODNs) were used as repair oligos, designed manually, and purchased from IDT. Sequences of sgRNAs and ssODNs can be found below. For gene editing, single cells were resuspended in BTXpress electroporation solution (VWR, Cat#732-1285), electroporated with 2 pulses of 65 mV for 20 ms in a 1 mm cuvette and transferred to 10 cm plates in StemFlex Medium (Thermo Fisher, Cat#A3349301) with Revitacell (RVC) supplement (1:100, Thermo Fisher, Cat#A2644501). Cells were selected with 350 ng/ml Puromycin for three days starting the day after electroporation. Edited single-cell clones were analyzed by RFLP assay, using enzymes HpyCH4IV (NEB, Cat#R0619S) for APP Arctic, and DdeI (NEB, Cat#R0175S) for APP Iberian, followed by Sanger sequencing as described^109^. Successfully edited clones were expanded and subjected to quality controls. The KI iPSC line is available from the authors after filling out an MTA.

### Quality controls of edited iPSC

Genomic integrity at the edited locus and the BCL2L1 gene was confirmed by quantitative genomic PCR as described previously^57,110^. Briefly, genomic DNA was extracted using the NucleoSpin Tissue Kit (Macherey Nagel, Cat#740952.250) and a RT-qPCR was performed using PrimeTime Gene Expression Master Mix (IDT, Cat#1055772) to determine copy numbers of the locus of interest in relation to a “house-keeping” locus (TERT gene). Undifferentiated state of iPSC was confirmed by immunostainings as described above using Oct4, Tra160, SSEA4, and Nanog antibodies. To verify the origin of each edited cell line, a microvariation (“fingerprint”) at the human D1S80 locus was analyzed by PCR. Off-target effects at the most probable loci, as predicted by CRISPOR, were excluded by PCR followed by Sanger sequencing. General genomic integrity was confirmed by molecular karyotyping using HumanOmni2.5Exome-8 BeadChip v1.4 (Life & Brain GmbH, Bonn).

### Amyloid-β (Aβ) measurements in supernatant

Media of 3-7 cultures (100 µl per culture) was conditioned for 5 days, collected on ice, spun down at 300 *g* at 4 °C for 5 min, transferred into protein-low-binding tubes (Eppendorf), flash frozen in liquid nitrogen and stored at -80 °C. For analysis, supernatants were thawed on ice, centrifuged at 11,000 *g* for 10 min at 4 °C, and used to measure Aβ38, Aβ40 and Aβ42 with the V-PLEX Plus Aβ Peptide Panel 1 Kit (6E10, Meso Scale Discovery, Cat#K15200G) according to manufacturer’s instructions. “Total Aβ” levels were calculated as the sum of the three different Aβ isoforms. Measured Aβ values were normalized to total protein or RNA levels of the cultures used for media conditioning.

### Sequential protein extraction

Protein from 3BTMs was extracted in increasingly harsh buffers, namely 1) DEA buffer (0,2% DEA in 50 mM NaCl, pH = 10), 2) RIPA buffer (20 mM TRIS-HCl pH 7.5, 15 mM NaCl, 1 mM Na_2_EDTA, 1 mM EGTA, 1% NP-40, 1% sodium deoxycholate, 2.5 mM pyrophosphate), and 3) Guanidine-based buffer (RP1, Macherey-Nagel, Cat#740934) as “insoluble” fraction. To minimize protein degradation during the isolation, all buffers were supplemented with protease inhibitor and all steps up to addition of the Guanidine buffer were performed on ice or at 4 °C.

For the first step, 20-25 3BTMs per sample were pooled, washed once with PBS, transferred into 100 µl DEA buffer, and dissociated with ceramic beads on a Precellys Evolution homogenizer as described above. The resulting cell suspension was spun down for 10 min at 2500 *g*, the supernatant transferred to an ultracentrifuge tube (Beckman Coulter) and centrifuged at 100,000 *g* for 30 min at 4 °C. The supernatant (“DEA fraction”) was neutralized by adding 0.5 M TRIS (pH = 6.75, 1:10), and flash frozen. The pellet was resuspended in 20 µl RIPA buffer, homogenized thoroughly together with the pellet formed after the initial centrifugation step, ultracentrifuged at 100,000 *g* for 30 min and the supernatant (“RIPA fraction”) flash frozen. The pellet was resuspended in 20 µl Guanidine buffer and spun down at 100,000 *g* for 30 min, and the resulting supernatant (“Guanidine fraction” = “insoluble fraction”) flash frozen. Protein concentrations of the Guanidine fraction (1:2 diluted) were measured using Pierce BCA Protein-Assay (Thermo Fisher, Cat#23227). Guanidine fractions were diluted 1:10 with Diluent 35 from the MSD Human Aβ V-PLEX kit (6E10), and samples were analyzed using the kit as above, and results normalized to total protein levels in the fraction.

### SDS-PAGE and Western Blotting

Total protein was extracted with RIPA buffer as described for mass spectrometry analysis. Proteins were separated on hand-casted 10% TRIS-Glycine gels (TGX Stain-Free FastCast Acrylamide Starter Kit, 10%, Bio-Rad, Cat#1610182), at 80 V for 25 min and at 120 V for 1-2 h, and transferred to nitrocellulose membranes (0.45 µm, VWR, Cat#10600002) at 100 V for 1 h on ice. Membranes were blocked in 3 % BSA in TBS-T (TBS with 0.1% Tween-20), incubated in primary antibody (AT270 1:400, PHF-1 1:200, K9JA 1:10,000) in 1 % BSA in TBS-T overnight at 4 °C, and washed 3x 10 min in TBS-T. Blots were then incubated in HRP-labeled secondary antibodies (1:10,000 in blocking solution) for 1 h at RT, washed 3x 10 min in TBS-T, and developed with Clarity Western ECL Substrate (BioRad, Cat#1705060) on a Fusion Fx7 imager (Vilber). Band intensities were measured in FiJi.

### Seeding of 3BTMs with Aβ42 and treatment with aducanumab or control antibody

For seeding experiments, lyophilized synthetic human Aβ42 (AggreSure beta-Amyloid (1-42), Hölzel Biotech, Cat#72216) was prepared as previously described^5^. In brief, Aβ42 was resuspended at 5 mM in DMSO and further diluted in PBS to a 100 µM stock solution. Resuspended peptide was incubated at 4 °C for 24 h to allow oligomerization before being frozen at -80 °C. For experiments, oligomerized synthetic Aβ42 was added into 3D culture maintenance media at 500 nM on days 8 and 10. Human aducanumab and non-binding control antibodies were obtained from Denali Therapeutics and added into 3D culture maintenance media at 1 µg/ml every 2-3 days starting on Day 20.

## Statistical analysis

No statistical methods were used to predetermine sample size, and the experiments were not randomized. “n” denotes the number of independent experiments, meaning cultures prepared from independent batches of cell differentiations were analyzed. In bar graphs, if not stated otherwise, each dot represents one experiment. Experimental data was analyzed for significance using GraphPad Prism v10.1 software unless stated otherwise. Multiplicity-adjusted p < 0.05 was considered statistically significant. Significance was analyzed either by 2-sided Student’s t-tests (paired or unpaired, as indicated) or Mann-Whitney test (for non-normal distribution of values as seen by Normality test in GraphPad), if only 2 groups were compared, one-way ANOVA if more than 2 groups were compared with 1 independent variable, two-way ANOVA if more than 2 groups were compared with 2 independent variables. Multiple-comparison post-testing with Tukey’s, Šidák’s or Dunn’s method was performed as recommended by GraphPad and indicated. Pearson’s correlation was calculated using GraphPad. Statistical details are described in the figure legends. All bar graphs are shown as mean + standard deviation (SD). Box plots show the median, 25^th^ and 75^th^ percentile (box), and 1.5x interquartile range (IQR) (whiskers).

## Data and code availability

The scRNA-seq profiling dataset has been deposited to the European Genome-phenome Archive (EGA) (EGAS50000000469), will be released as of the day of publication and made accessible upon reasonable request to the Data Access Committee (DAC). The mass spectrometry proteomic data have been deposited to the ProteomeXchange Consortium via the PRIDE partner repository with the dataset identifier PXD055254 and are publicly available as of the day of publication.

Original western blot images and microscopy data reported in this paper will be shared by the lead contact upon request.

All original code to analyze the scRNAseq data as well as tools/packages information to fully reproduce the analysis will be made publicly available on GitLab as of the date of publication. Any additional information required to reanalyze the data reported in this paper is available from the lead contact upon reasonable request.

## Extended Data Figures

**Extended Data Fig 1.**
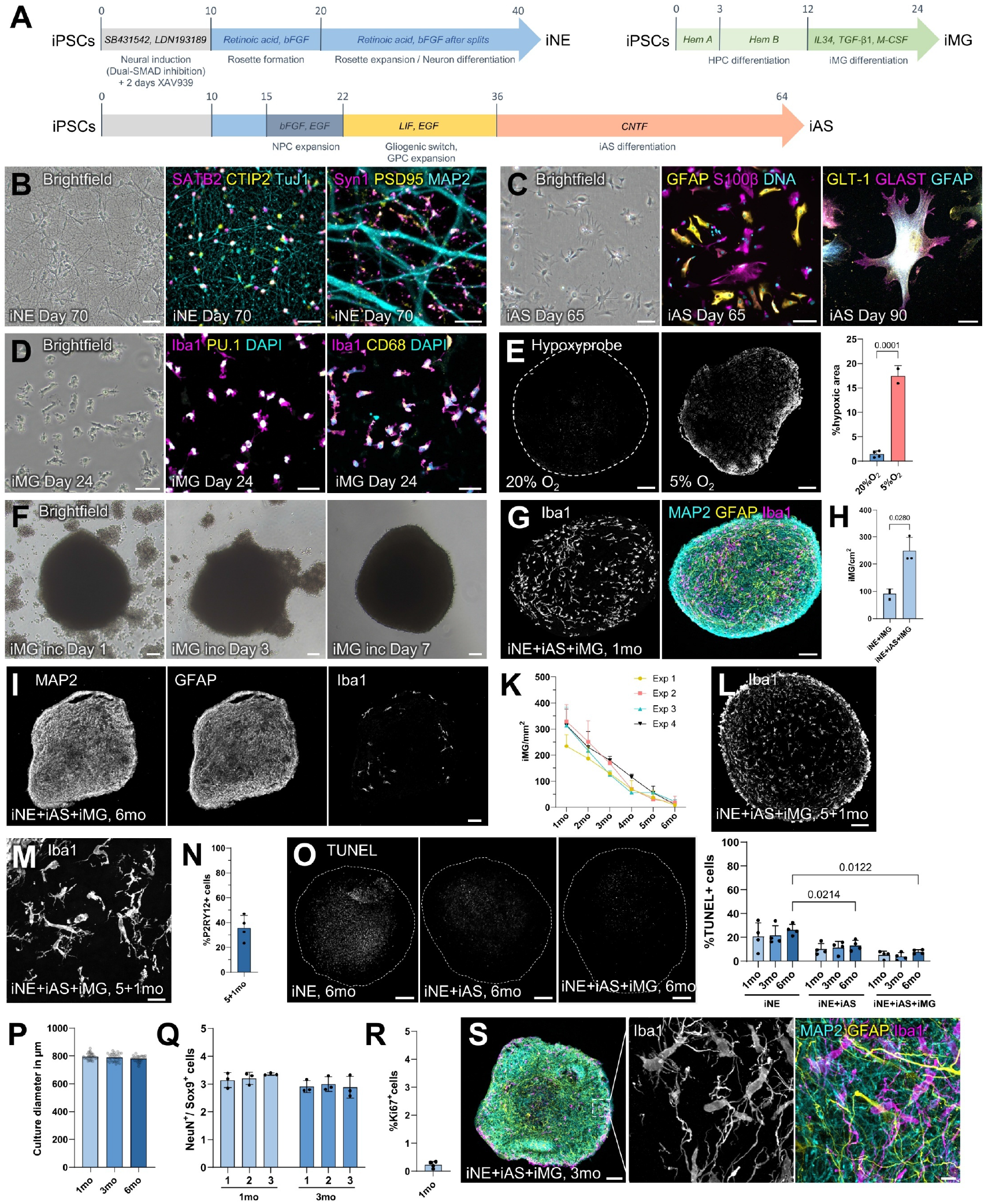
Differentiation of iPSC-derived brain cell types and general characterization of 3BTMs. **(A)** Overview of protocols used to differentiate induced pluripotent stem cells (iPSCs) into cortical neurons (iNE), astrocytes (iAS), and microglia (iMG). **(B)** iNE in 2D monoculture on Day 70 after iPSC stage showing typical morphologies and marker expression. Left: Brightfield image, scale bar 50 µm. Middle: IF staining showing expression of cortical markers SATB2 and CTIP2 and beta-III tubulin (TuJ1), scale bar 50 µm. Right: IF staining showing synapse formation of iNE by colocalization of presynaptic Synapsin-1 (Syn1) and postsynaptic PSD95 on MAP2-positive neurites, scale bar 5 µm. **(C)** iAS in 2D monoculture on Day 65 showing typical star-shaped morphologies and marker expression. Left: Brightfield image, scale bar 100 µm. Middle: IF stainings for S100beta and GFAP (scale bar 100 µm). Right: IF stainings for GLT-1 and GLAST (scale bars 20 µm). **(D)** iMG in 2D monoculture on Day 24 showing typical morphologies and marker expression. Left: Brightfield image, scale bar 50 µm. Middle: IF stainings for Iba1 and PU.1, scale bar 30 µm. Right: IF stainings for Iba1 and CD68, scale bar 30 µm. **(E)** IF stainings in 3-month-old iNE+iAS 3BTM using Hypoxyprobe, to label proteins in hypoxic cells^96^, under normal (20% O_2_) and hypoxic conditions (5% O_2_, positive control). Quantification on the right (n = 2-4 experiments with two 3BTMs each, each dot represents one experiment, unpaired two-sided t-test), scale bars 100 µm. **(F)** Brightfield images showing migration of iMG towards and into 3BTM over the course of one week, scale bars 100 µm. **(G)** IF staining of iNE+iAS+iMG 3BTM at 1 month of age for Iba1, MAP2 and GFAP, scale bars 100 µm. **(H)** Quantification of iMG numbers in 3D co-culture without and with iAS at 1 month (n = 3 experiments with two 3BTMs each, each dot represents one experiment, paired two-sided *t*-test). **(I)** IF staining of 3BTM at 6 months showing the different cell types, scale bar 100 µm. **(K)** Quantification of iMG numbers in 3BTMs over time (n = 4 experiments with two 3BTMs each, represented by colored lines). **(L)** IF staining of 6-month-old iNE+iAS+iMG culture repopulated with iMG at 5 months, scale bar 100 µm. **(M)** Close-up of IF staining as in (M), scale bar 20 µm. **(N)** Quantification of the percentage of P2RY12-positive cells at 6 months in cultures repopulated with iMG at 5 months of age. **(O)** IF TUNEL stainings of 3BTMs containing only iNE, iNE+iAS, or iNE+iAS+iMG at 6 months, scale bars 100 µm. Quantification on the right (n = 4 experiments with two 3BTMs each, each dot represents one experiment, Repeated measures two-way ANOVA with Tukey’s multiple comparisons test, data for 3 months timepoint also shown in Figure 1L). **(P-S)** Confirmation of 3BTM properties in an independent iPSC line. **(P)** Quantification of culture diameters at 1, 3 and 6 months in an independent iPSC line, each dot represents one culture, (n = 3 experiments with 10-11 3BTMs per experiment and timepoint, each dot represents one 3BTM). **(Q)** Quantification of iNE:iAS ratio at 1 and 3 months in an independent iPSC line (n = 3 experiments with two 3BTMs per experiment and timepoint, each dot represents one experiment). **(R)** Quantification of Ki67-positive cells in an independent iPSC line at 1 month (n = 3 experiments with one 3BTM each, each dot represents one experiment). **(S)** IF staining of a 3-month-old iNE+iAS+iMG culture of an independent iPSC line, scale bar 100 µm. Zoom-in on the right, scale bar 10 µm. Data are represented as mean + SD.

**Extended Data Fig 2.**
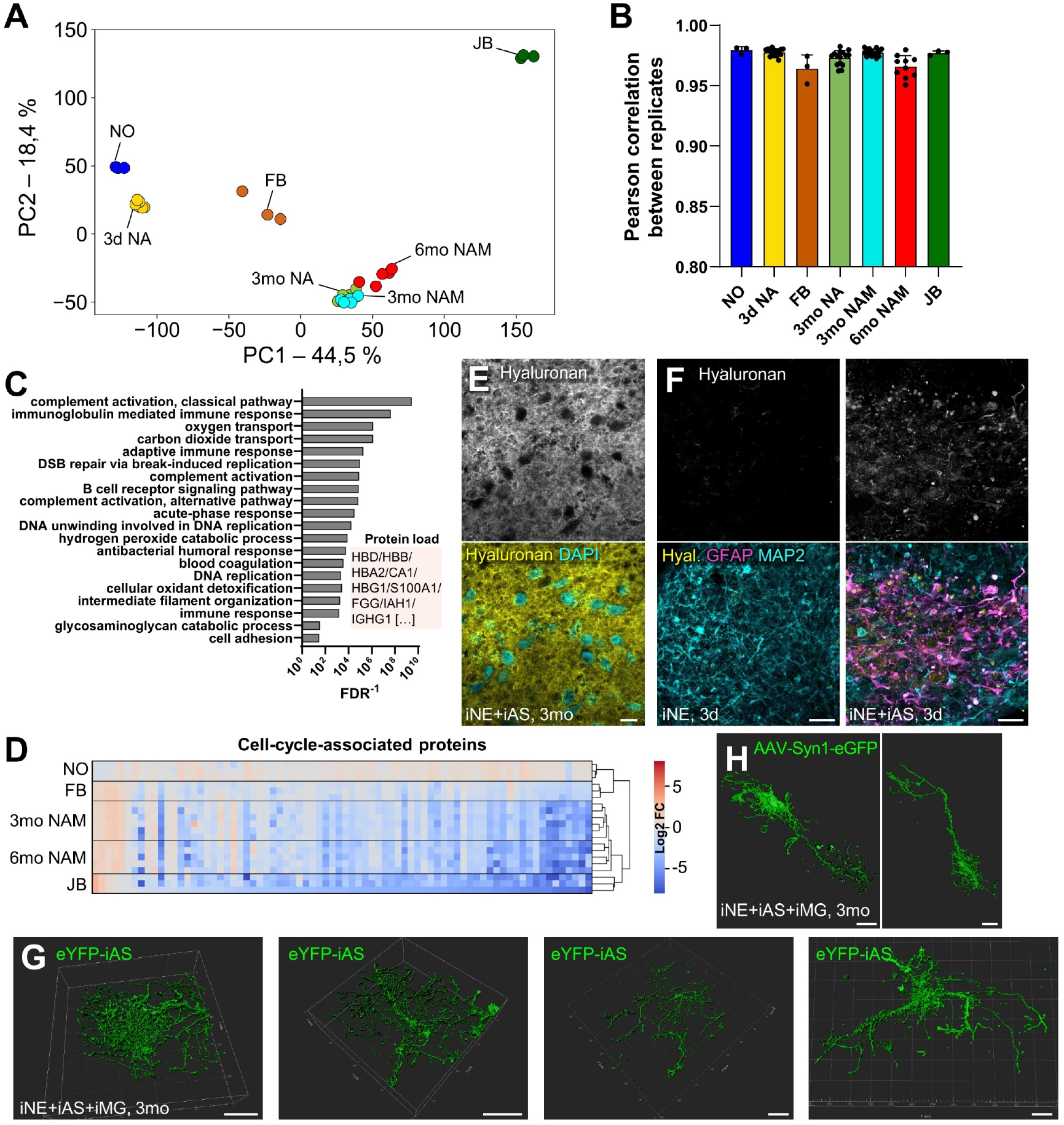
Mass spectrometry analysis of 3BTMs and single-cell morphologies. **(A)** Principal component analysis (PCA) of proteomic dataset based on all proteins after imputation showing PC1 and PC2. **(B)** Pearson correlation of log2 LFQ intensities of all detected proteins between replicates of each sample Data presented as mean + SD. **(C)** Gene ontology analysis (GO-BP) of top proteins influencing PC2, sorted by false-discovery rate (FDR). Note that PC2 is largely driven by “blood-dependent” processes, such as complement activation, carbon dioxide/oxygen transport, or B cell receptor signaling, as well as DNA replication. A subset of the top proteins contributing to PC2 load is shown. **(D)** Heatmap showing log2-fold changes (log2FC) in protein levels of cell-cycle-associated proteins found in proteomics dataset compared to 3-day-old iNE+iAS cultures (set to zero). Hierarchical clustering of the different samples on the right. **(E)** IF staining showing extracellular deposition of hyaluronan, as seen by exclusion of staining in DAPI-positive cell bodies, scale bar 10 µm. **(F)** IF staining for hyaluronan in 3BTMs containing only iNE or iNE+iAS at 3 days, scale bars 20 µm. **(G)** 3D rendering of eYFP-labelled, single iAS, scale bars 50 µm. **(H)** 3D rendering of AAV-hSyn1-eGFP-labelled, single iNE, scale bars 20 µm.

**Extended Data Fig 3.**
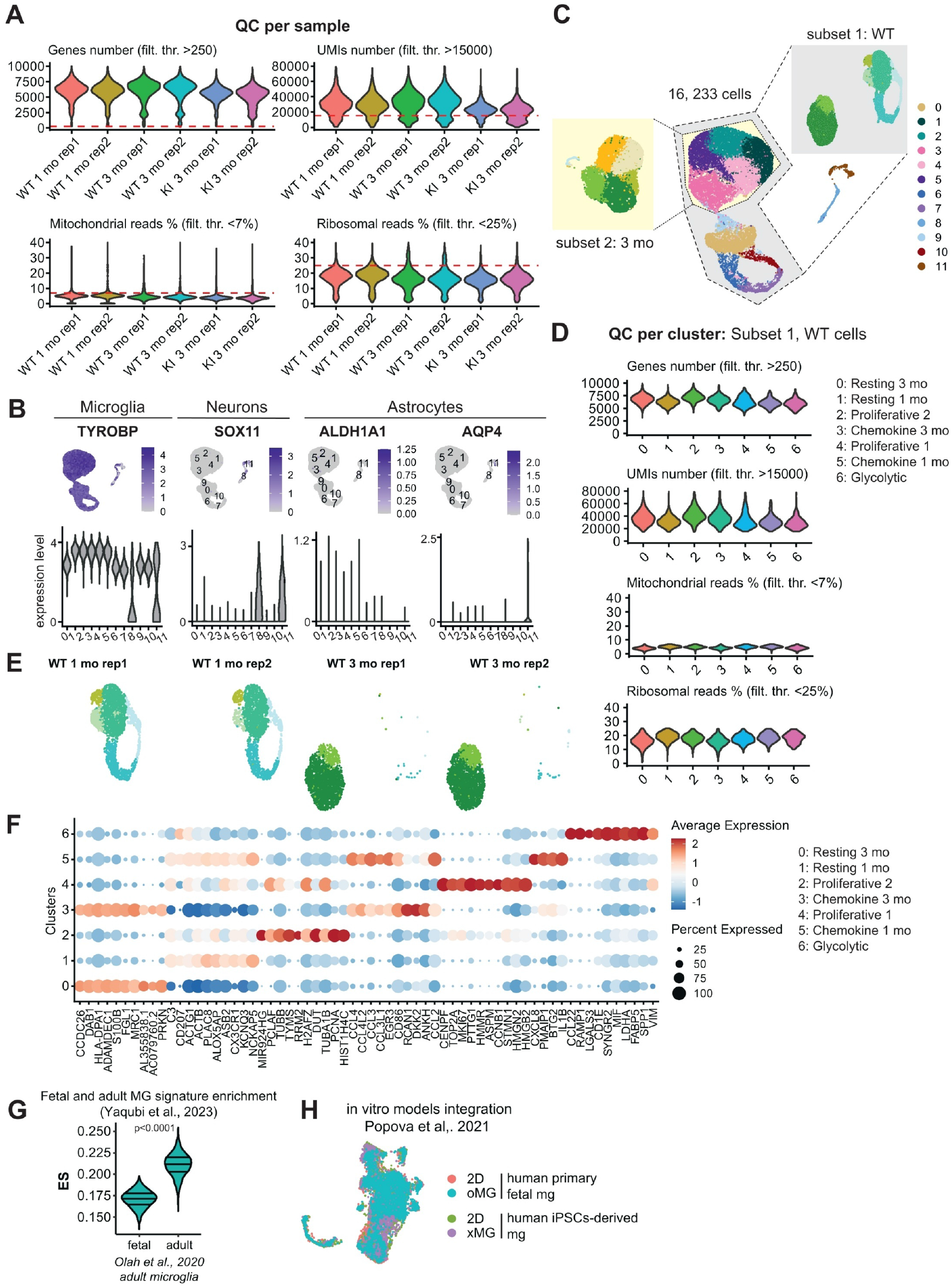
Single-cell RNA sequencing analysis of iMG from 3BTMs at 1 and 3 months. **(A)** Per cell quality controls (QC) across all clusters including counts per cell (>250), UMIs per cell (>15,000), mitochondrial reads (<7% per cell) and ribosomal reads (<25% per cell). **(B)** UMAP-based overview of the overall dataset subsetting strategy: clusters 8 and 11 were excluded from the initially QC-filtered dataset, considering their expression of neuronal and astrocytes markers. WT (Figure 3) and 3mo (Figure 6) subsets were produced for downstream analyses. **(C)** Expression of markers for microglia (*TYROBP*), neurons (*SOX11*) and astrocytes (*ALDH1A1, AQP4*) across the identified dataset clusters. Only pure iMG clusters were selected for downstream analyses (clusters 8 and 11 were excluded). **(D)** Per cell QC as in (A) across the WT subset clusters (Figure 3B). **(E)** UMAP reported in Figure 3B split by biological replicates. **(F)** Top 10 marker genes (only positive, detected in >25% of cells with log_2_FC>0.25, ordered by log_2_FC) across the WT subset clusters (Figure 3B). **(G)** Violin plot reporting fetal and adult MG signature ES in adult human MG (dataset by Olah et al.^46^) used as external dataset for signature validation. **(H)** UMAP representation of the integrated Popova et al.^21^ *in vitro* model dataset, including primary human fetal MG cultured in 2D (2D, human primary fetal) or incorporated into cortical organoids (oMG, human primary fetal) as well as human iMG either cultured in 2D (2D, human iPSCs-derived) or xenotransplanted into mouse brain (xMG, human iPSCs-derived).

**Extended Data Fig 4.**
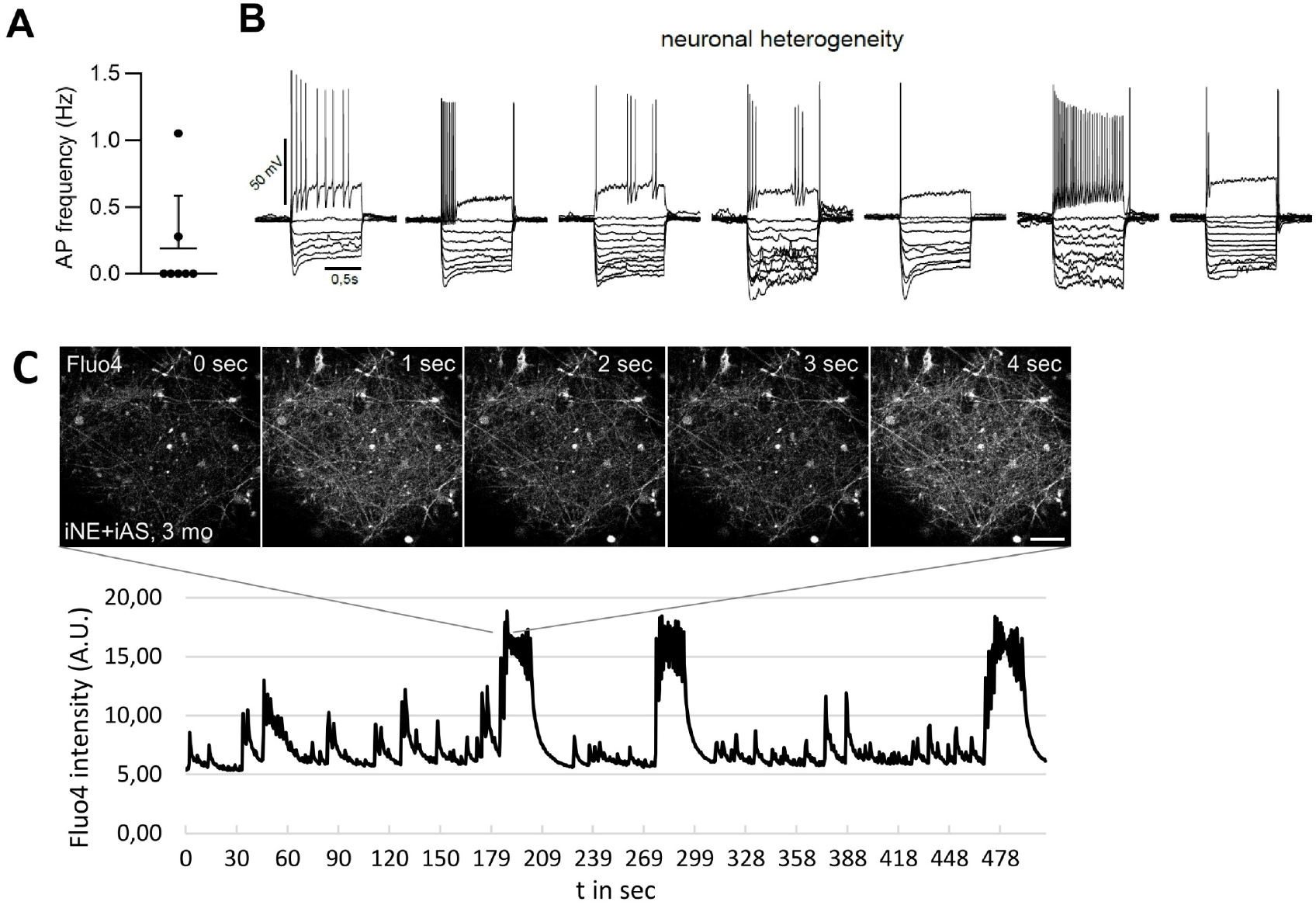
Functional assays for iNE. **(A)** Spontaneous action potential (AP) frequency observed in patch-clamp measurements of single iNE in iNE+iAS+iMG 3BTMs at 3 months (n = 2 experiments with 3-4 cells measured from one 3BTM each, each dot represents one cell). **(B)** Evoked APs observed as in (A) showing heterogeneous firing patterns in response to depolarizing stimuli. **(C)** Top: Example images of live cell calcium imaging using Fluo-4 in iNE+iAS+iMG 3BTMs at 3 months showing spontaneous and synchronized activity. Bottom: Quantification of Fluo-4 intensity over time, timeframe shown above is marked by lines.

**Extended Data Fig 5.**
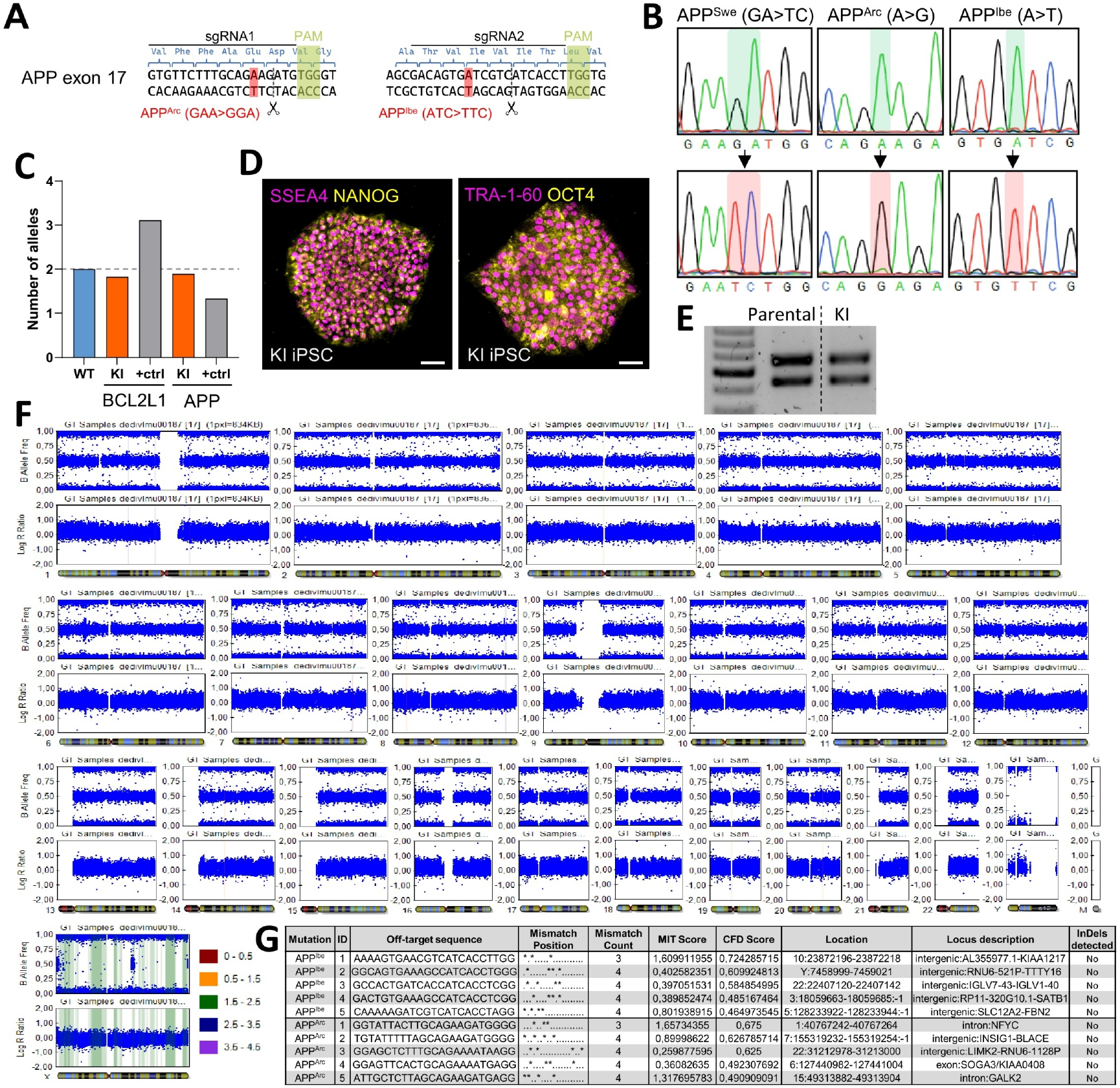
Gene editing and quality controls to introduce AD-causing APP mutations. **(A)** Overview of the edited loci in APP exon 17, sgRNAs used for editing are marked by black solid lines, associated protospacer adjacent motives (PAM) marked in green, Cas9 cut sites marked by dashed black line and scissors, mutations to be inserted are shown in red (Arctic = APP^Arc^, Iberian = APP^Ibe^). **(B)** Sanger sequencing confirming presence of the APP Swedish (APP^Swe^) mutation and successful editing for APP^Arc^ and APP^Ibe^. **(C)** Quantitative genomic PCRs (qgPCRs) showing normal copy number of 2 alleles at *BCL2L1* locus compared to a positive control with 3 alleles, and at the edited locus (*APP* exon 17) compared to positive control with 1 allele. **(D)** IF stainings showing expression of markers for undifferentiated state SSEA4 and NANOG (left) as well as TRA-1-60 and OCT4 (right), scale bars 50 µm. **(E)** Fingerprinting PCR showing descent of edited KI lines from the parental line by same band pattern. **(F)** Molecular karyotyping of KI cell line to analyze integrity of all chromosomes after editing. **(G)** Overview of most-likely off-target loci (according to crispor.tefor.net). Each locus was analyzed by Sanger sequencing and no off-target editing was found.

**Extended Data Fig 6.**
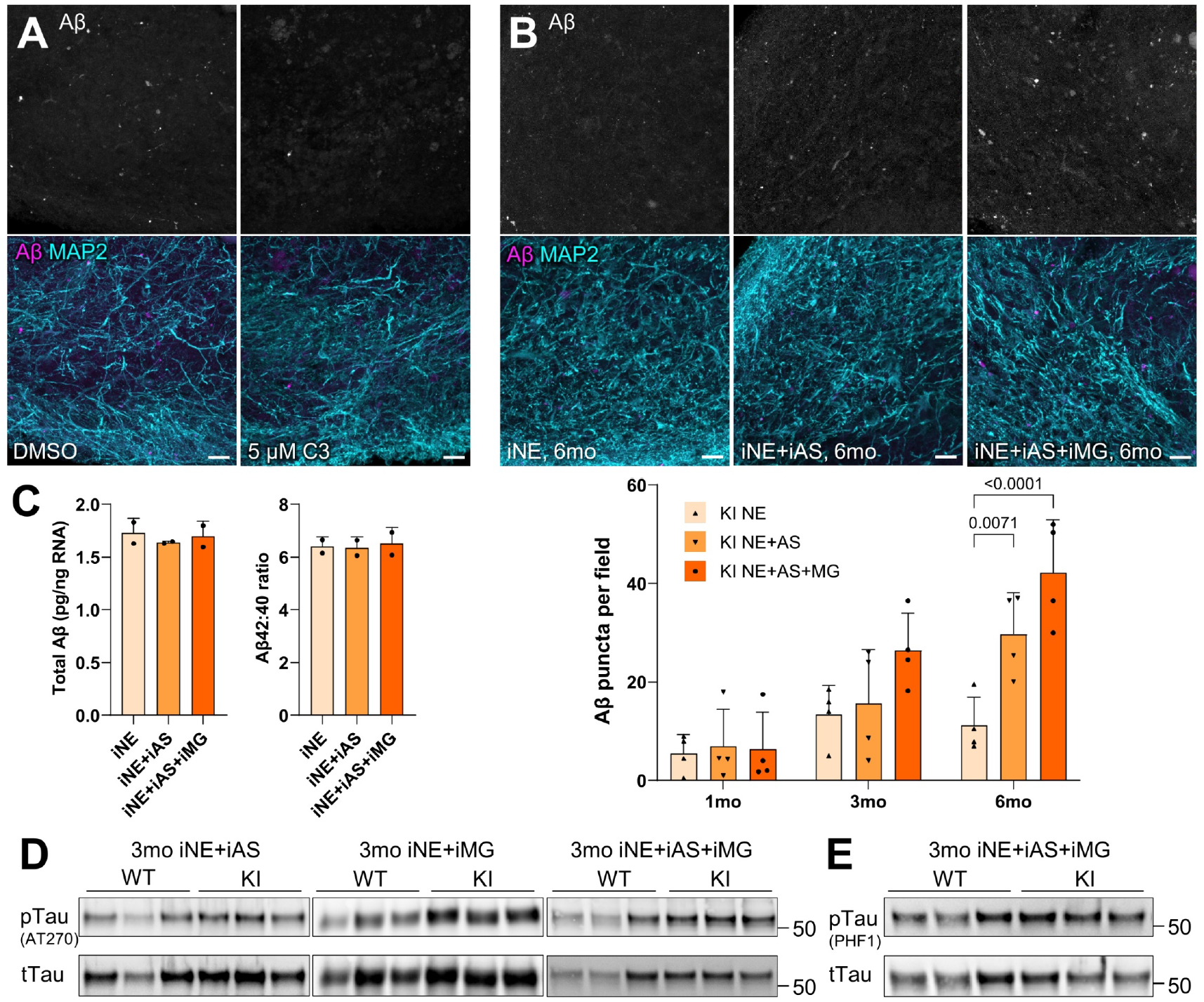
Gene editing and quality controls to introduce AD-causing APP mutations. **(A)** IF stainings of KI 3BTMs at 6 months treated with DMSO (loading control), or 5 µM C3 (BACE-inhibitor IV) showing reduced Aβ accumulation after C3 treatment, scale bars 20 µm, quantification in Figure 5E. **(B)** IF stainings of KI 3BTMs with different cell type combinations at 6 months, scale bars 20 µm. Quantification of Aβ puncta on the bottom showing a time course for each combination (n = 4 experiments with 1-2 3BTMs each, each dot represents one experiment, two-way ANOVA with Tukey’s multiple comparisons test). **(C)** Immunoassay to detect Aβ in supernatants of 1-month-old KI 3BTMs containing different cell type combinations showing total Aβ levels (left) and Aβ42:40 ratio (right) (n = 2 experiments with 4-5 3BTMs used for media conditioning each, each dot represents one experiment). **(D)** Western blot (WB) analysis for phospho-Tau (pTau, AT270 antibody – Thr181 epitope) and total Tau (tTau, K9JA antibody) of 3-months-old KI 3BTMs containing different cell types, quantification in Fig. 5G, 5H. **(E)** WB analysis as in (D) for pTau using the PHF1 antibody (pSer396/pSer404), quantification in Fig. 5I. Data are presented as mean + SD.

**Extended Data Fig 7.**
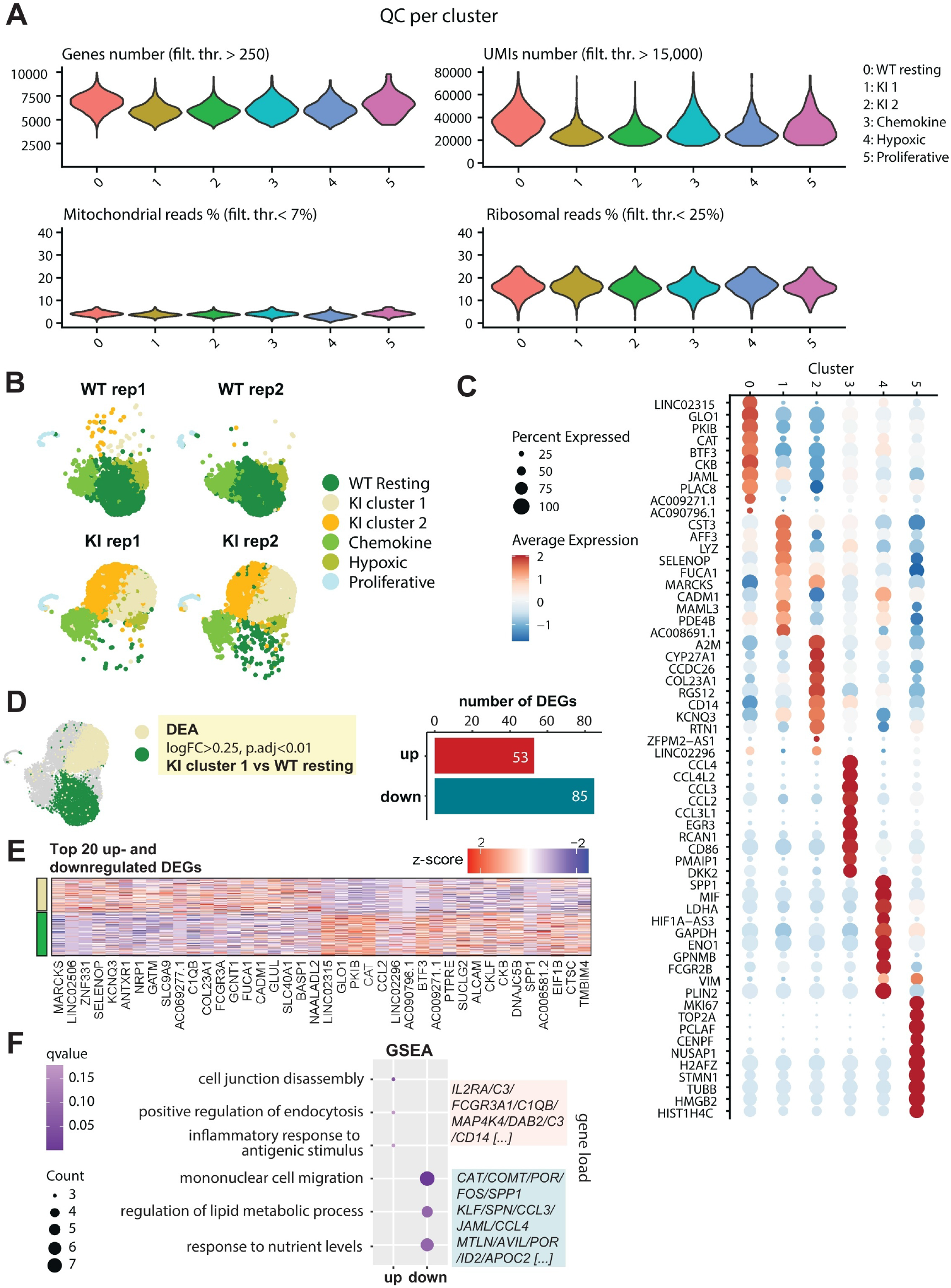
Single-cell RNA sequencing analysis of WT and KI iMG at 3 months. **(A)** Per cell quality controls across all WT and KI clusters at 3 months (Figure 6C), including counts per cell (>250), UMIs per cell (>15,000), mitochondrial reads (<7% per cell), and ribosomal reads (<25% per cell). **(B)** UMAP reported in Figure 6C split by biological replicates. **(C)** Top 10 marker genes (only positive, detected in >25% of cells with log_2_FC>0.25, ordered by log_2_FC) for each cluster. **(D)** Number of differentially expressed (DE) genes (p.adj<0.01, logFC>0.25) obtained by comparing “*KI cluster 1*” and “*WT resting*” clusters. **(E)** Heatmap showing normalized scaled expression of the top 20 up- and downregulated DEGs. **(F)** Gene Set Enrichment Analysis (GSEA) terms enriched in the up- and down-regulated genes resulting from “*KI cluster 1*” vs “*WT resting*” clusters DE comparison (*q*-value<0.2). A list of interesting up- and downregulated genes contributing to the load of GSEA functional terms is shown.

